# Chinese hamster tissue and CHO cell transcriptomes reveal immortalization signatures and the factors driving lineage divergence

**DOI:** 10.64898/2026.11.25.769185

**Authors:** Debabrata Mohanty, Deepikka Senthilkumar, Prahalaad Vijay Varahaswami, Nikshep Grampurohit, Da Eun Kang, Rajeev Ranjan, P.M. Meera Alagu Sundari, Matthew Myint, Jong Kwang Hong, Kuin Tian Pang, Meiyappan Lakshmanan

**Author notes:** Correspondence to: M. Lakshmanan. These authors have equally contributed to the work (Debabrata Mohanty, Deepikka Senthilkumar).

## Abstract

Chinese Hamster Ovary (CHO) cells dominate biotherapeutic manufacturing but still have a few limitations in post-translational modifications such as N-glycan branching, O-glycan core extension and inefficient protein folding/processing. It is yet unclear whether these CHO-specific traits reflect general Chinese hamster biology or arise specifically from cell line immortalization. We assembled a uniformly processed transcriptome compendium of 350 RNA-seq samples from 24 studies, integrating nine native Chinese hamster tissues with four CHO lineages (CHO-K1, CHO-S, CHO-DG44, CHO-DXB11). While CHO cells retain core proliferative and biosynthetic machinery when compared to Hamster tissues, it has undergone coordinated and selective loss of tissue-specific functions, and thus establishing these are immortalization-driven traits rather than general hamster-biology origin. Focused analysis of CHO cell lines further revealed that parental lineage and culture media as the dominant drivers of observed transcriptional heterogeneity where the cell identity is responsible for transcriptome variation in a higher proportion of genes. By integrating genome copy number or alternate promoter usage with gene expression data, we observed that while gene copy number variations contributions are almost negligible, alternate promoter usage governs the variation in about 20% of genes that showed cell line specific patterns while 60% had a buffering effect, indicating that additional, yet to be characterized regulatory layers drive CHO cell heterogeneity. Together, these findings establish a genomic and regulatory framework for understanding CHO lineage divergence, with direct implications for rational cell line selection and engineering in biotherapeutic manufacturing.

**Graphical abstract:** 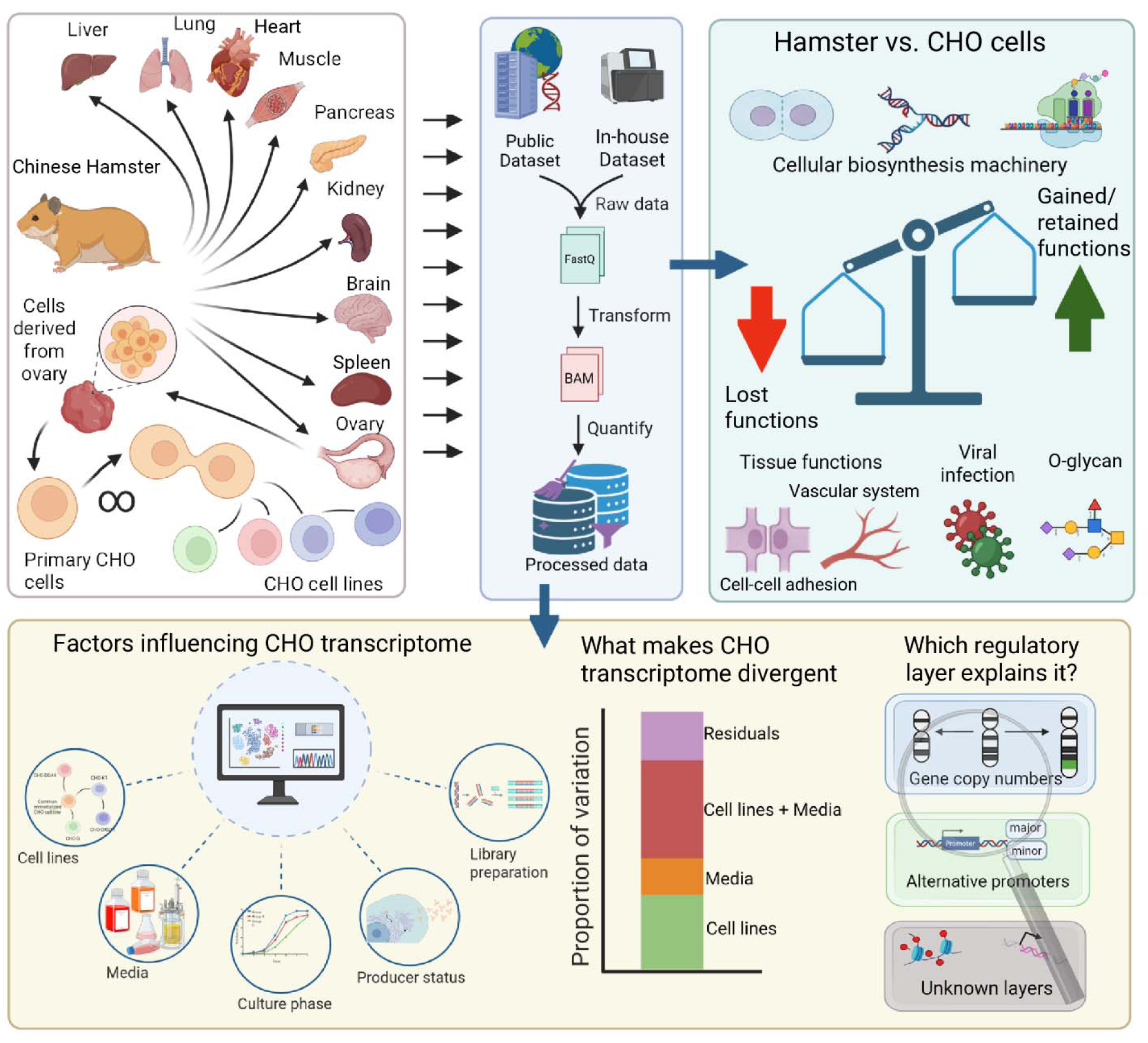

## Introduction

Chinese Hamster Ovary (CHO) cells are regarded as the primary mammalian host choice for recombinant therapeutic protein production due to their several unique properties, but particularly due to their ability to make human-like post-translational modifications including N-glycosylation (Lakshmanan et al. 2019). Since their isolation from the Chinese Hamster in 1956 by Dr. Theodore Puck, CHO cells have been routinely cultivated in cell culture and extensively used for biotherapeutic manufacturing (Reinhart et al. 2019). The continuous cultivation of CHO cells has led to multiple mutations and genetic and epigenetic drifts, either induced intentionally or unintentionally, resulting in multiple, phenotypically distinct cell lines (CHO-DG44, CHO-DXB11, CHO-K1 and CHO-S), which have actually diverged from a common ancestor over decades (Wurm 2013). The variations in genetic make-up contribute to the differences in phenotypes when these parental CHO cells are applied in biotherapeutic production (Reinhart et al. 2019; Lakshmanan et al. 2019; Széliová et al. 2020; Z. Hu et al. 2013), yet the regulatory basis of this cell line divergence remains poorly resolved. Beyond the high industrial relevance, the lineage divergence of CHO cells offers an excellent model system where we can dissect how genomic, transcriptional, and regulatory layers jointly manifest cellular phenotypes under sustained, artificial selective pressure, a broader question of interest in genome biology.

To understand the impact of genetic and molecular variations on the phenotype, omics technologies have been widely used to systematically characterize how intricate molecular and cellular interactions manifest complex phenotypic changes. Whole genome transcriptome analysis is a primary tool for characterizing the cell line and process specific differences due to its ability to investigate the expression of individual gene products across conditions, as well as its simplicity in the methodology and low-cost requirement when compared to other methods. CHO transcriptome datasets are now available across a multitude of conditions including the investigation of differences between a producer and non-producer, low-and high-producer, mAb vs. Epo-Fc producer, temperature shift during bioprocess, serum-free media adaptation, clonal variations and epigenetic regulation of gene expression, among others. Being a prevalent omic approach, a large number of transcriptomics dataset are now available in the public repositories such as the Sequence Read Archive (SRA) and the European Nucleotide Archive (ENA), which can be compiled and uniformly processed to establish a large-scale transcriptome compendium for a systematic analysis of gene expression profiles across various culture conditions and different CHO cell lines.

Previously, several attempts have been made to build a CHO transcriptome compendium over the past decade. Rupp et al., 2014 developed the first publicly usable annotated CHO-K1 transcriptome using both reference-and *de novo*-assembly pipelines and made it accessible to the scientific community (Rupp et al. 2014). Another study compiled RNA-Seq and microarray data of Chinese hamster tissue samples and CHO cell lines to compare gene expression profile across different cell lines and between tissues and cell lines, focusing on glycolysis and glycosylation-related expression differences and the contribution of SNVs (Vishwanathan et al. 2015). Later, an online CHO transcriptome compendium consisting of 23 different transcriptome datasets from previous studies was built and analyzed cell line specific variation across CHO-S, CHO-K1 and CHO-DG44 cell lines and identified the key transcriptomic signatures among these cell lines giving rise to various cell line specific traits (Singh et al. 2018). In a recent effort, a significantly larger transcriptome compendium with more than 640 RNA-seq datasets, consisting of three CHO cell lines CHO-K1, CHO-DXB11 and CHO-S was assembled and analyzed it over two different published works (Riedl et al. 2025, 2026). They showed that while gene expression largely varies based on the cell lineages, a subset of the genes showed variations based on experimental conditions, keeping a core set of transcripts constantly expressed in all the cell lines. While these studies collectively demonstrated the utility of large-scale transcriptome analysis, two fundamental gaps remain unresolved. First, whether cell-line-specific traits such as lack of bisecting GlcNAc and increased viral resistance is applicable to Chinese Hamsters in general or just observed at the immortalized CHO cell lines. Second, and more fundamentally, none of the studies have systematically analyzed which factors, whether genomic dosage, cis-regulatory promoter activity or other mechanisms, drives the cell-line-specific transcriptional divergence observed in CHO cell lines despite the availability of large-scale datasets.

Herein, we address both the abovementioned gaps by first building a publicly accessible, the most comprehensive CHO transcriptome compendium, comprising of RNA seq datasets from 9 hamster tissues (brain, liver, lung, kidney, ovary, spleen, pancreas, heart, muscle) and four commonly used CHO cell lines, CHO-K1, CHO-S, CHO-DG44 and CHO-DXB11. Using this resource, we first analyze whether commonly reported CHO cell-line specific traits in literature such as increased viral resistance are due to immortalization by comparing the cell line specific transcriptome with hamster tissues. We show that CHO cells retain core proliferative and biosynthetic machinery while undergoing coordinated and selective loss of tissue-specific functions such as cell adhesion and vascular signaling. We then compared the gene expression profile of different CHO cell lines and reasoned whether the observed transcriptional variation is due to genomic dosage (copy number variation) or cis-regulatory changes (promoter usage), revealing that neither fully accounts for observed divergence indicating the as-yet-uncharacterized regulatory mechanisms as key drivers of CHO cell line heterogeneity. Together, these findings establish a genomic and regulatory framework for understanding CHO cell line divergence, with direct implications for rational cell line selection and engineering in biotherapeutic manufacturing.

## Results

### Development of the Chinese Hamster transcriptome compendium

We developed the comprehensive Chinese Hamster transcriptome by first surveying the list of available RNA-seq data from public repositories Sequence Read Archive (SRA) and European Nucleotide Archive (ENA). We focused on studies which either used Chinese Hamster tissues or different CHO cell lines grown in a vast array of conditions and shortlisted 23 unique Bioprojects. Note that we also included the RNA-seq data available from in-house Chinese Hamster tissue samples. The compendium has a total of 350 RNA-seq samples from 24 different studies, comprising of 9 Hamster tissues (brain, heart, kidney, liver, lung, muscle, ovary, pancreas and spleen) and 4 different CHO cell lines (CHO-DG44, CHO-DXB11, CHO-K1 and CHO-S) from 2 and 22 different studies, respectively **(Supplementary file 1, Table 1).** It should be noted that the CHO cell lines either express a recombinant protein such as mAb or just the parental cell lines grown in a variety of conditions such as adherent and suspension cultures in batch or fed-batch with various serum-containing or serum-free media. We downloaded raw data, i.e., FASTQ files, of all the samples from the repositories and uniformly processed them using a standardized RNA-seq analysis pipeline to obtain the gene expression counts (**Figure 1a**). At the outset, we noted that each of the datasets followed diverse library preparation methods and had different sequencing depth (**Figure 1b**) and varying sequence lengths (**Figure 1c**). Following the uniform processing, we extracted the counts of protein coding genes and further normalized them to remove any potential batch effects that may arise from the differences in library preparation for sequencing, library layouts and sequencing platform. The processed is accessible through a web interface (<u>CHO_transcriptome</u>).

**Fig 1.**
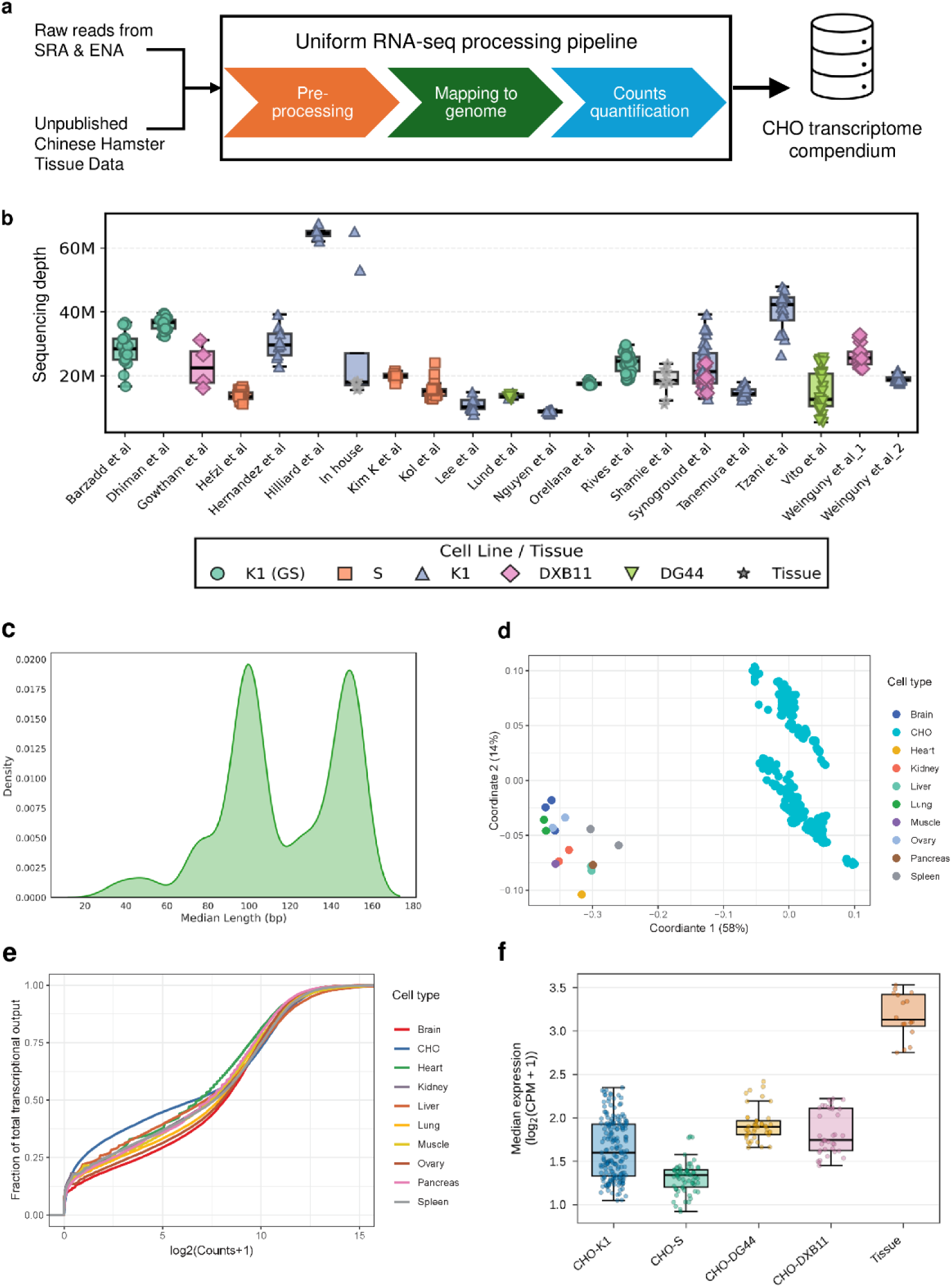
Overview of the assembled CHO transcriptome compendium. (a) Schematic of the compendium workflow, in which raw reads from SRA & ENA and unpublished Chinese hamster tissue data were processed through a uniform RNA-seq pipeline (pre-processing, mapping to genome, count quantification) to generate the CHO transcriptome compendium. (b) Sequencing depth of samples acros studies included in the compendium, grouped by cell line (K1 (GS), S, K1, DXB11, DG44) and Chinese hamster tissue. (c) Density distribution of median read length (bp) across samples. (d) Multi-dimensional scaling of samples based on gene expression profiles, coloured by cell type, showing separation of CHO cell lines from Chinese hamster tissues. (e) Cumulative distribution of the fraction of total transcriptional output as a function of log2(Counts+1), shown for each cell type. (f) Median gene expression (log2(CPM+1)) of samples from CHO-K1, CHO-S, CHO-DG44, CHO-DXB11 and Chinese Hamster tissues.

On average, we found the expression of 14066±1107 genes out of the total 21841 protein-coding genes in the Chinese Hamster across all samples. Interestingly, we found 357 (∼1.6%) genes showed a very low expression, i.e., almost equal to zero, across all the Hamster tissues as well as CHO cell line samples analyzed **(Supplementary file 2, Table 1)**. When we further analyzed the role of these 357 genes, we noted that among the enriched pathways, most of the genes are contributing to sensory perception of smell (GO:0007608; *adj. P* < 0.05) and Odorant binding (GO:0005549; *adj. P* < 0.01). Since our dataset collection does not have samples from nasal tissues, it is unsurprising that the expression of these genes was not found across the entire dataset. Similarly, we also noted sensory perception of taste (GO:0050912; *adj. P* < 0.05) was also enriched in this set of genes which did not have any expression in the assembled dataset as it does not involve any samples related to tongue tissues **(Supplementary file 2, Table 2)**.

Next, we analyzed the global gene expression patterns of the expressed genes (at least in one of the conditions) using multi-dimensional scaling (MDS). It highlighted that the expression patterns of the CHO cell lines were markedly different from the 9 hamster tissues, including the ovary tissue from which the CHO cells were originally isolated (**Figure 1d**). We also compared the transcriptional output of the nine tissues and CHO cells and found that CHO cells express fewer genes and show lower expression levels for several genes (**Figure 1 e-f**).

### Coordinated loss of tissue-specific functions and gain of repressors expression highlight selective transcriptional rewiring in CHO cells

To further comprehensively understand the gene expression differences between CHO cells and hamster tissues, we performed differential expression analysis and found three categories of genes: 1) expressed in CHO cells but not in hamster tissues (4 genes), 2) expressed in hamster but not in CHO cells (464 genes) and 3) reduced expression levels in CHO cells when compared to hamster tissues (3077 genes) (**Figure 2a**) **(Supplementary file 2, Tables 3-6).**

**Fig 2.**
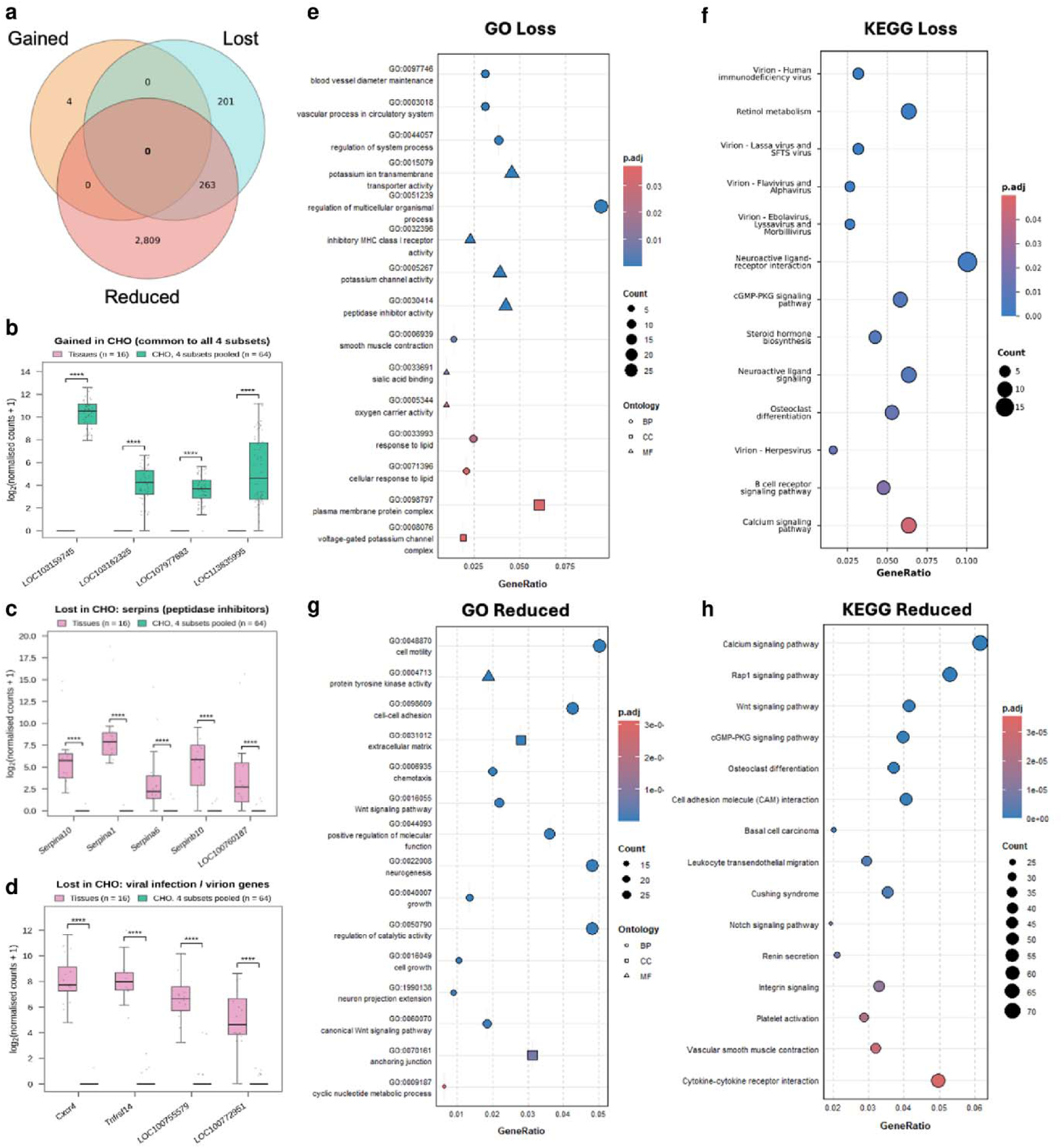
Functional enrichment analysis of differentially expressed genes in CHO cell lines relative to Chinese Hamster tissues. (a) Venn diagram of genes gained, lost and reduced in CHO cells relative to hamster tissues. (b) Expression of genes gained in CHO cells that are common to all four subsets in tissues and CHO cells. (c) Expression of serpins (peptidase inhibitors) lost in CHO cells. (d) Expression of viral infection / virion genes lost in CHO cells e) GO enrichment analysis of DEGs showing loss of expression in CHO cell lines relative to different types of Chinese hamster tissues. f) KEGG pathway enrichment analysis of DEGs showing loss of expression in CHO cell lines relative to different types of Chinese hamster tissues g) GO enrichment analysis of DEGs showing reduced expression in CHO cell lines relative to different types of Chinese hamster tissues h) KEGG pathway enrichment of DEGs showing reduced expression in CHO cell lines relative to different types of Chinese hamster tissues.

Interestingly, all the 4 genes which have gained expression in CHO cells have a DNA binding activity (**Figure 2b**). Particularly, *LOC103162325* and *LOC107977683* are zinc finger proteins containing Krüppel associated box (KRAB) domain (Moosmann et al. 1997), where the former gene is reported to have repressor activity for RNA pol II dependent transcription and the later has repressor activity for both RNA pol I and RNA pol II mediated transcription (Wang et al. 2021). Another gene which gained function as compared to tissues is *LOC103159745*, whose mouse ortholog is reported to have transcription repression function affecting the regulatory region of RNA pol II. Collectively, the gain of expression of these three genes which are reported to have transcription repression activity point towards a potential role in the reduction of overall gene expression in CHO cell lines.

Next, we analyzed the 464 genes that have lost expression in CHO cells when compared with Hamster tissues. Multiple genes with endopeptidase inhibitor activity including *Serpina10*, *Serpina1*, *Serpina6* and *Serpinb10* were found to have lost expression in CHO cells (**Figure 2c**). Since endopeptidase inhibitors are essential for maintaining extracellular matrix homeostasis and systemic balance of immune responses, CHO cells could have selectively lost the expression of these genes as they neither have the necessity to maintain the ECM environment nor have inflammatory responses. We also found that genes from response to lipid (GO:0033993; adj. *P* < 0.05) and cellular response to lipid (GO:0071396; adj. *P* < 0.05) processes have also lost expression where most of the genes are involved in response to lipopolysaccharide, i.e., response to bacterial infection. This suggests that as CHO cells are maintained in sterile conditions for extended periods, it may have led to their progressive loss in expression. Similarly, expression of genes involved in the responses to viral infections were also found to be lost in CHO cells as compared to tissues (**Figure 2d**). Particularly, genes encoding for cell surface receptors of multiple viruses such as HIV, Flavivirus, Ebola virus, Lassa virus, Alpha virus and Herpes virus were all lost the expression, confirming that CHO cells have reduced susceptibility for viral contamination. It should be further highlighted that most of the GO terms enriched for the genes that have lost expression in CHO cells belong to tissue level function rather than core cellular function (**Figure 2e & f**). For example, processes related to circulatory systems such as vascular processes in the circulatory system (GO:0003018; adj. *P* < 0.01) and blood vessel diameter maintenance (GO:0097746; adj. *P* < 0.01), as well as liver specific molecular function such as haptoglobin binding (GO:0031720; adj. *P* < 0.01) and oxygen carrier activity (GO:0005344; adj. *P* < 0.01) were enriched. Interestingly, three genes from sialic acid binding (GO:0033691; adj. *P* < 0.01) (*LOC100751896*, *LOC100753971* and *LOC100764017*), belonging to immunoglobulin lectin family and involved in several immune responses, have lost expression in CHO cells. Together, these results suggest that continuous *in vitro* cultivation could have led to loss of complex functional pathways specific to tissue characteristics due to the lack of interaction with other cells and absence of architectural complexity in the tissue microenvironment. (Pathway enrichment file is available in **supplementary file 2, Table 11**)

Similar to genes that have lost expression as compared to tissues, the genes that have reduced expression in CHO cells were also evaluated. A total of 3077 genes were found to have reduced expression as compared to nine hamster tissues. Further functional analysis revealed pathways related to cell adhesion are enriched in all the three GO terms (**Figure 2g**). Similarly, KEGG enrichment analysis also revealed genes from various pathways related to cellular adhesion such as cell adhesion molecules, Notch signaling pathway, Focal adhesion, ECM-receptor interaction, Tight junction, Adherens junction was enriched (**Figure 2g**; see **Supplementary file 2, Table 12**). These observations together highlight that CHO cells have reduced cell-cell interaction and cell-matrix interaction as compared to Hamster tissues. We also found that *Wnt* signalling pathways to be significantly enriched in both biological process (GO:0016055; adj *P* < 0.01) and molecular function (GO:0017147; adj *P* < 0.05) of GO terms, and KEGG pathways (cge04310; adj *P* < 0.01). Here, it should be noted that though *Wnt* signalling pathway is involved in multiple functions such as cell cell adhesion, focal adhesion and membrane trafficking (Tejeda-Muñoz and Mei 2024; Rask et al. 2003), the set of genes differentially expressed in Hamster vs CHO cells comparison from *Wnt* signaling pathway were all related to cell fate determination and signaling pathways regulating pluripotency of stem cells, suggesting they are involved primarily in developmental process. Notably, their expression is not completely lost, which may be due to their evolutionarily conserved functional roles (Yu et al. 2024) and could also be attributed to their ovarian origin where the role of *Wnt* signalling is indispensable (Hernandez Gifford 2015).

Several genes related to protein glycosylation have also reduced expression in CHO cells when compared to hamster tissues. For example, Mucin type O-glycan biosynthesis was significantly enriched (cge00512; adj. *P* < 0.05) where multiple N-acetylegalactosamine transferases (*Galnt12*, *Galnt16*, *Galnt17*, *Galnt18*, *Galnt3* and *Galnt6*) have reduced expression. Similarly, *C1galt1c1l*, *B3gnt3* and *Gcnt4* genes, which are required for core 1 and core 2 biosynthesis (Ho et al. 2016) have also reduced expression in CHO cells as observed earlier (Yang et al. 2014; Riedl et al. 2025; Xu et al. 2011). We also noted multiple genes with Hexosyltransferase activity (GO:0016758; adj *P* < 0.01), which are important for synthesis of type - I chain of N-glycan and O-glycans (Holgersson and Löfling 2006), are found to have reduced expression in CHO cells, similar to the observation in previous studies (Cherian et al. 2015). Further, *B3galt5*, *B3gnt3*, *B3gnt4* and *B3gnt7*, all of which are required for mucin type O-glycan core-4 extension (Ho et al. 2016) also show reduced expression in CHO cells. Additionally, we also noted *Mgat1* and *Mgat3* to have reduced expression in CHO cells when compared tissues. It should be noted that *Mgat3* is essential for forming bisecting GlcNAc structure and has already been reported to have reduced expression in CHO cells (Vishwanathan et al. 2015). Since multiple glycosylation related genes have their gene expression reduced in CHO cells, we further investigated whether protein processing and secretory pathways are affected in the CHO cells when compared to Hamster tissues. For this purpose, we mapped the genes involved in post translational protein processing pathways using SecRecon as it systematically categorizes the genes by their specific functional role in the secretory machinery, e.g., ER quality control and vesicular trafficking at different levels (Masson et al. 2025). 4 out of the total 5 systems in secRecon (post translational modification, vesicle trafficking, proteostasis and protein conformation) were significantly enriched (**Supplementary file 2, Table 14**). Among multiple subsystems within post translational modification, Glycosphingolipid (adj. *P* < 0.01) and O-glycosylation (adj. *P* < 0.01) showed statistically significant enrichment (**Figure 3a**). Notably, multiple genes mapped to the O-glycosylation subsystem from secRecon overlapped with the KEGG pathway O-glycan biosynthesis, e.g., *Galnt16*, *Galnt17*, *Galnt18*, *Galnt3*, *Galnt6*, *Lfng*, *Mfng*, *Ogt*, *Pomt2*. Similarly, *A4galt, Abo, B3galt2* and *St3gal6* genes which are majorly involved in glycosphingolipid biosynthesis were also enriched (**Figure 3b**). Within the post translational modifications system, even though other subsystems such as N-glycosylation, glycosaminoglycans, glycerophospholipids, and phosphorylation were also enriched, although they were not statistically significant. Together, these observations indicate that the glycosylation repertoire of CHO cells could be narrower than the tissues.

**Fig 3.**
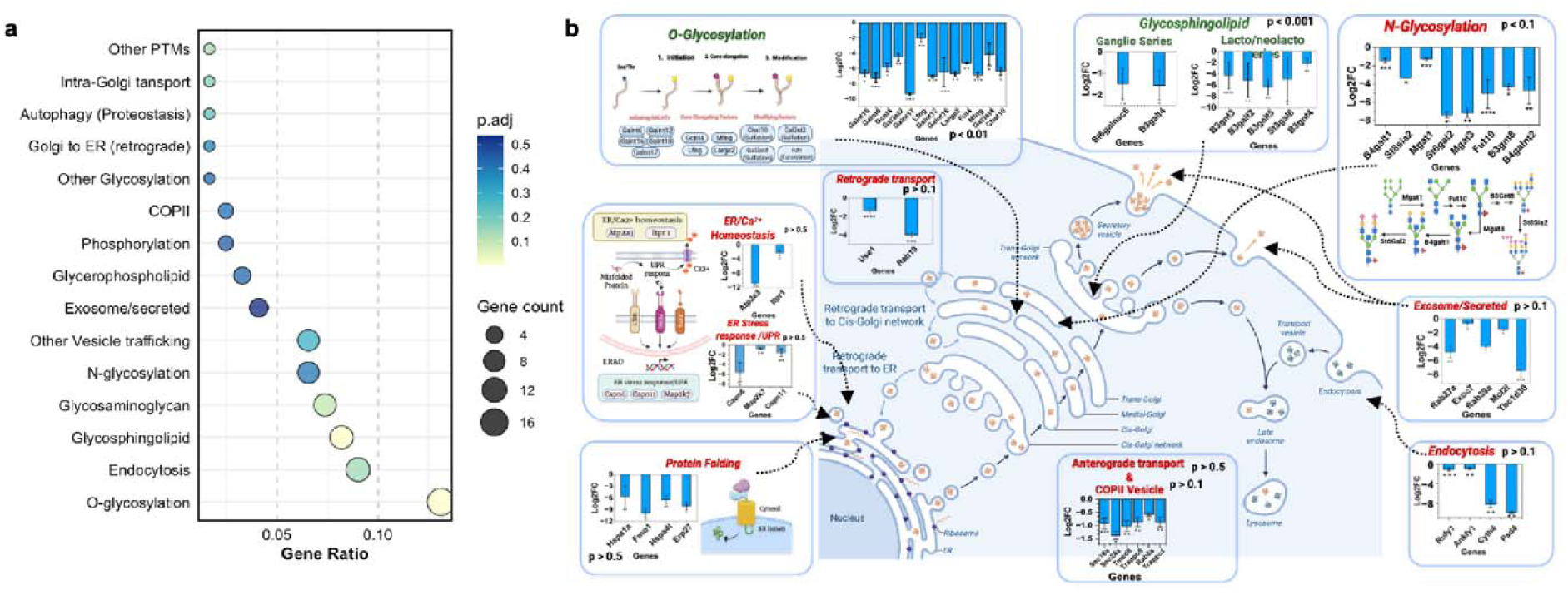
Functional enrichment of key secretory pathway components downregulated in CHO cell lines relative to Chinese Hamster tissues. (a) Bubble plots displaying the enrichment of downregulated/reduced expression of genes mapped to the secretory pathway via SecRecon model at the subsystem level. (b) Schematic overview of the secretory pathway delineating identified systems and subsystems, annotated with representative genes and their corresponding subcellular site of action. P-values were adjusted using Benjamini - Hochberg correction and P-value < 0.05 was considered as statistically significant.

Overall, our comparison between CHO cells and Chinese Hamster tissues indicates that while CHO cells preserve the cellular characteristics by maintaining the gene expression required for important cellular functions such as cell division and DNA replication and translation, but have selectively lost the expression of genes that are responsible for tissue characteristics. Notably, these observations are highly in concordance with the earlier study which analyzed the proteomic differences between CHO cells and hamster tissues (Heffner et al. 2020). Moreover, our analysis also shows that genes involved in post translational modifications and viral infection are either lost or reduced in expression as their corresponding pathways are enriched in both categories, corroborating with the known observation that CHO cells display intrinsic resistance against viral infections (Berting et al. 2010; Chiang et al. 2019).

### Lineage background and culture media are primary drivers of CHO cell lines transcriptome variation

We next shifted our focus towards analysing the variation within CHO cell lines as the compiled dataset involved four different parental CHO cell lines (CHO-K1, -DXB11, -DG44 and -S) and their producer clones (producing various recombinant proteins including IgG, erythropoietin, etc.) grown in different media and modes of cultivation (adherent vs. suspension, batch vs. fed-batch), and the samples were harvested at different phases of cell culture. The global analysis of the CHO cells revealed a characteristic transcriptional signature, as captured by UMAP, where a cell line specific grouping was observed: while CHO-S and CHO-DG44 clustered separately and individually from other cell lines, CHO-K1 and CHO-DXB11 clustered together (**Figure 4a**). Here, it should be noted that these observations are in line with known genetic background of the CHO parental cell lines, where CHO-K1 and -DXB11 arise from same lineage, while the other two cell lines are from different lineages, although all of them are derived from a common ancestor (Lewis et al. 2013) (**Figure 4b**). In addition to cell lines, culture media type was also found to have some contribution towards the observed groups. Therefore, we used linear models to understand how each of the variables, i.e. cell line, media, growth phase and producer/non-producer, contributes to the overall variation in gene expression patterns among the total genes which are expressed in at least one of the conditions. If the observed gene expression variation is not explained using any of these variables, then the model fits it as residues. Notably, the observed variation in the ∼16% (3304/20774) of expressed genes could not be explained by any of the fitted variables as the residuals were the top contributor (≥ 70% of observed variation) (**Figure 4c**). Among the fitted variables, cell line background was observed to be the top contributor for gene expression variations - a total of 314 genes (∼2%) showed a dominant contribution from cell line variation (≥ 70% of observed variation) (Supplementary file 2, Table 15). Genes such as *Msh3*, *Pon2*, *Ctps2*, *Hes2*, *B4galt6*, *Ptp4a3*, *Kitlg* and *Ebf3* showed only cell line specific expression variation (≥ 95% of observed variation) where *Hes2* is expressed only in CHO-DG44 while *Pon2* and *Ptp4a3* are expressed only in CHO-DG44 and CHO-S, *Ctps2* and *Kitlg* are expressed only in CHO-DXB11 and CHO-K1 (**Figure 4d**). Second, a set of 132 genes (∼0.5%) showed gene expression variation based on the cell culture media (≥ 70% of observed variation). For example, *Ephb2* and *Pou3f4* showed higher expression only in ActiPro (Hyclone) and CHO-S-SFMII (Gibco) media, respectively (**Figure 4e**). Moreover, we observed a minor set of genes (<0.1%) such as *Taf7l*, *Defb124*, *Fam3d*, *Mgp*, *Prrg3* and *Svip*, showed growth phase specific variations (≥ 70% of observed variation), while no specific set of genes showed appreciable variation between producer and non-producer cells (maximum variation captured <30% only). Apart from a specific list of genes which explained the variance entirely based on cell line or media, we also noted a set of 3298 genes (∼16%) that showed a cumulative contribution from both cell line and media (≥ 70% of observed variation). We performed pathway enrichment analysis of these genes and noted that these two variables have effects on pathways related to cellular survival and stress response and structural remodeling **(Supplementary file 2, Table 16)**. In addition, pathways related to translation (cge03010; Ribosome - adj. *P* < 0.01) and cellular adhesion (cge04520; Adherens junction and cge04510; Focal adhesion - adj. *P* < 0.01 in both) were also enriched in this category. This indicates that both culture media type and CHO cell line heterogeneity together exert broad and systemic effects that reprogram fundamental cellular regulatory networks, resulting in functional variation in the phenotypes.

**Fig 4.**
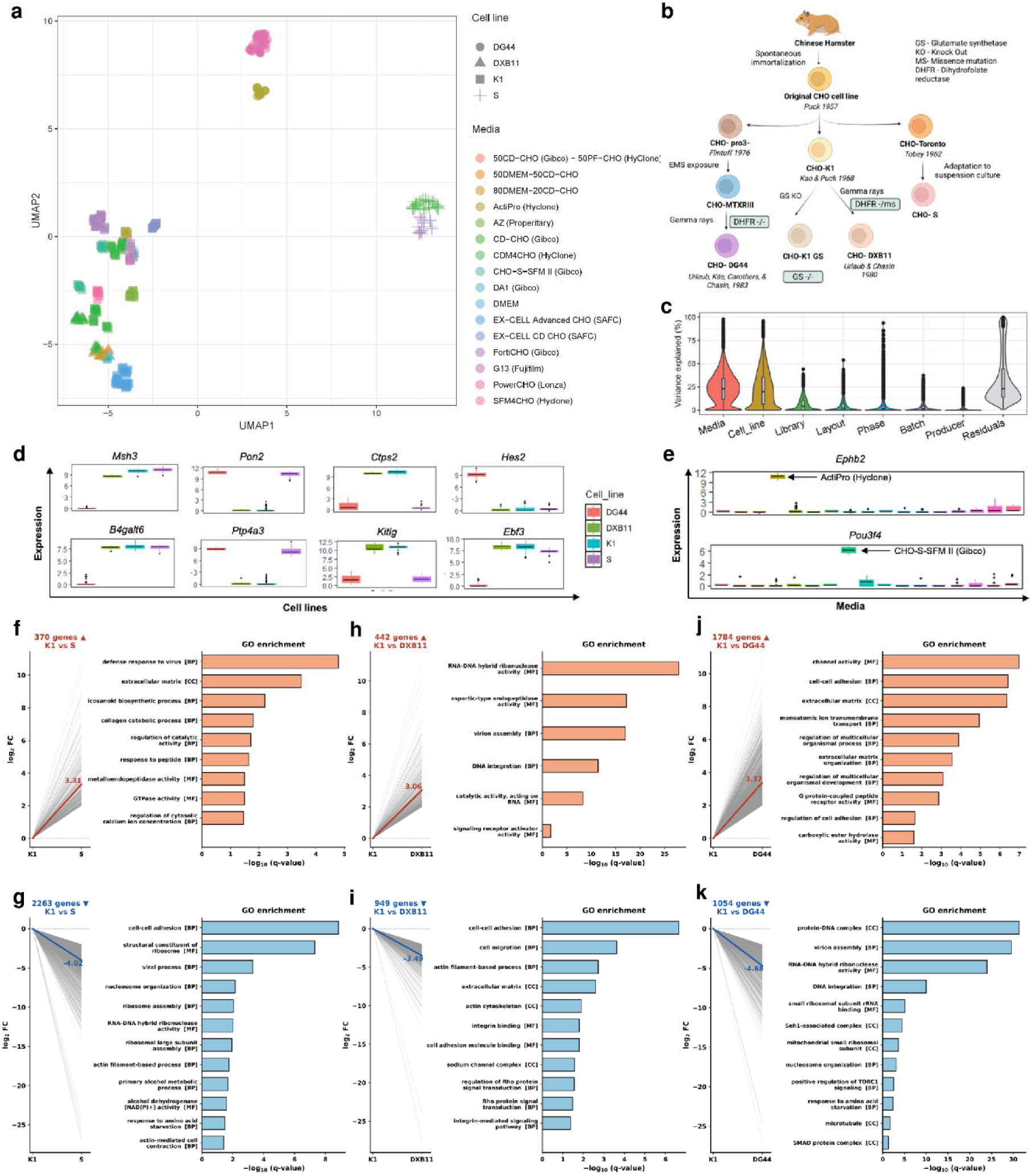
Lineage background and culture media are the primary drivers of CHO transcriptome variation. (a) UMAP of CHO samples based on gene expression, coloured by culture medium and shaped by cell line. (b) Lineage of the CHO parental cell lines (CHO-K1, CHO-DXB11, CHO-DG44, CHO-S) derived from the original CHO cell line. (c) Proportion of variance in gene expression explained by each variable (variancePartition). (d) Expression of genes whose variation is dominated by cell lines (Msh3, Pon2, Ctps2, Hes2, B4galt6, Ptp4a3, Kitlg, Ebf3). (e) Expression of genes whose variation i dominated by culture medium (Ephb2 in ActiPro; Pou3f4 in CHO-S-SFM II). (f–k) Differentially expressed genes relative to CHO-K1 and their GO enrichment for CHO-S (f, g), CHO-DXB11 (h, i) and CHO-DG44 (j, k); f, h, j show upregulated genes and g, i, k show downregulated genes. P-values were adjusted using Benjamini - Hochberg correction and P-value < 0.05 was considered as statistically significant.

As parental cell lines were one of the major drivers of CHO transcriptome heterogeneity, we analyzed the key gene expression differences among the CHO cell lines using CHO-K1 as a reference. CHO-DG44 and CHO-S showed almost the same number of differentially expressed genes (2838 and 2633) while CHO-DXB11 showed only half of that number (1391) **(Supplementary file 2, Tables 17-19)**. Enrichment analysis of the differentially expressed genes (DEGs) identified that both CHO-S and CHO-DG44 showed a large number of pathways both in GO terms and KEGG as compared to CHO-DXB11 (**Figure 4 f-k, Supplementary file 2, Tables 20-25**). Downregulation of amino acid starvation was commonly observed in CHO-S and CHO-DG44. Particularly, *Seh1*, one of the critical components of the GATOR2 complex (Chantranupong et al. 2014), is downregulated in both CHO-S and CHO-DG44. GATOR2 plays an important role in keeping the mTOR pathway active by inhibiting GATOR1 (Takahara et al. 2020), while its reduced expression may suppress the downstream functions of the mTOR signalling pathway. Moreover, ribosome assembly (GO:0042255, adj *P* < 0.01 in CHO-S), structural constituent of ribosome (GO:0003735: adj *P* < 0.01 in both CHO-S and CHO-DG44) and mTOR signaling pathway (adj *P* < 0.01 in CHO-DG44) and positive regulation of TORC1 signaling (adj *P* < 0.01 in CHO-DG44) were also found to be downregulated in CHO-S and CHO-DG44. Similarly, apoptosis related pathways such as negative regulation of apoptotic process (GO:0043066, adj *P* < 0.05 in CHO-S) and Apoptosis (cge04210, adj *P* < 0.01 in CHO-DG44) were also enriched in these two cell lines. These pathways are reported to be affected as a response to amino acid limitation (Yatipanthalawa et al. 2026) and collectively, it indicates that CHO-S and CHO-DG44 are more sensitive to amino acid starvation, which may affect their anabolic processes and promote cell death. Pathways having significant effect on epigenetic regulation were also enriched in both CHO-S and CHO-DG44. For example, nucleosome organization (GO:0034728: adj *P* < 0.01 in both CHO-S & CHO-DG44), nucleosome assembly (GO:0006334: adj *P* < 0.01 in both CHO-S & CHO-DG44), structural constituent of chromatin (GO:0030527: adj *P* < 0.01 in both CHO-S & CHO-DG44) and ATP-dependent chromatin remodeling (cge03082: adj *P* < 0.05 in both CHO-S & CHO-DG44) were all downregulated, suggesting the potential epigenetic regulation of gene expression in these two CHO cell lines.

Interestingly, cell-cell adhesion and cell-matrix adhesion related pathways were downregulated in CHO-S and CHO-DXB11, while an increased expression was observed for CHO-DG44. Several cadherin protein coding genes such as *Cdh9*, *Cdh13* and *Cdh7* have shown reduced expression in CHO-S while *Cdh11*, *Cdh8* and *Cdh3* along with few claudin genes (*Cldn23*, *Cldn1* and *Cldn16*) show increased expression in CHO-DG44. Upregulation of these pathways suggests the tendency of CHO-DG44 towards cell-cell adhesion and aggregate formation during culture despite having anti-clumping agents in the media as cell adhesion molecules are reported to increase cell-cell aggregation in suspension culture (Malm et al. 2020; Louie et al. 2020). Another interesting observation is the upregulation of pathways related to actin filament in CHO-DG44, while the similar pathways were downregulated in other two cell lines. Besides maintaining the cytoskeletal structure of the cell, actin filaments have a significant role in protein secretion. Particularly, *Myo1g*, *Myo1a* and *Actc1* which play crucial roles in intracellular trafficking were all upregulated in CHO-DG44.

Apart from cell adhesion related pathways, several genes from cellular transport and metabolic pathways were upregulated, indicating that CHO-DG44 could have higher metabolic activity than CHO-K1. To test this hypothesis, we mapped the metabolic genes with *i*CHO3K, a comprehensive genome-scale metabolic network of Chinese Hamster, resulting in mapping of 2582 genes. Among the three comparisons, CHO-DG44 has the highest number of metabolic genes upregulated (223), while CHO-S has the largest number of downregulated genes (258) **(Supplementary file 2, Table 26)**. We also noted that a set of extracellular solute carrier transporters and fatty acid metabolism genes were upregulated in CHO-DG44 while it is downregulated in CHO-S. Unlike CHO-S and CHO-DG44, CHO-DXB11 has a comparatively lesser number of enriched pathways indicating the transcriptional similarity with CHO-K1, which could be possible because as both CHO-K1 and CHO-DXB11 arises from the same genetic lineage. Collectively, the cell line specific comparison identified CHO-DG44 to be metabolically more active than CHO-K1, CHO-S and CHO-DXB11 while identifying the sensitivity to amino acid deprivation in CHO-S and CHO-DG44. CHO-DXB11 remains functionally comparable to CHO-K1, highlighting the influence of evolutionary lineage on the cellular phenotype and functional behaviour. This is in line with the previous studies which emphasizes the cell line lineage to be the major determinant of the physiological behavior of the CHO cell lines (Riedl et al. 2025; Singh et al. 2018).

Since our analysis indicated media to be a significant contributor to the observed variation in transcriptome, we further analyzed the effect of media within CHO-K1 cells. CD-CHO (Gibco) medium was used as reference as it is one of the older baseline chemically defined formulations and frequently used as a standard reference chemically defined media in academic and industrial benchmarking studies. Relative to CD-CHO, a large number of genes were differentially expressed across the other media. The highest number of upregulated genes were present for the G13 (2100) followed by CHO-S-SFM II (2088) while the highest number of downregulated pathways were observed in EXCELL-Advanced CHO (4952) medium followed by FortiCHO (4294) and EXCELL CD CHO (3403) **(Supplementary file 3, Table 1-9)**. Pathway enrichment analysis also mirrored this trend, where the highest number of pathways were upregulated in G13 medium. In general, the enriched pathways among differentially expressed genes were mostly related to metabolic processes, transport activities along with cell-cell communication pathways (**Supplementary file 3, Table 10-18**) while downregulated pathways were related to epigenetic regulation. It should be highlighted that these media-specific signatures are consistent with known mechanisms by which medium composition shapes cell state. The relative changes in concentrations of nutrients such as amino acid, glucose and lipid regulate the cellular metabolism while osmolality and ionic composition variations differentially modulate transporter and adhesion programs. Additionally, the variations in growth factors and hydrolysate compositions alter signaling and cell-surface interactions while concentrations of certain key substrates or cofactors may also influence chromatin state and the expression of epigenetic regulators. Notably, unlike many other media, CDM4CHO showed least differentially expressed genes, indicating the transcriptional profile is highly similar to that of CD-CHO. However, this contrasts with an earlier report which showed that CDM4CHO supports better cell proliferation and mAb production than CD-CHO (Rodrigues et al. 2012), suggesting that improved culture performance does not necessarily require extensive transcriptome remodeling and may instead arise from post-transcriptional or metabolic differences.

### Copy number variations and genomic dosage are not primary drivers of CHO cell transcriptomic heterogeneity

It has been earlier reported that copy number variation (CNV) of certain genes drives the hard-wired transcriptome changes across CHO cell lines. Therefore, we next analyzed CNV variation among the cell lines by keeping CHO-K1 as reference. Among the three cell lines, CHO-DG44 showed the highest fraction of genes with CNV, i.e., 6.13%, while CHO-S had 3.64% and CHO-DXB11 showed 2.1% (**Supplementary file 4**). The highest number of genes with copy number loss were observed in CHO-DG44 followed by CHO-S and CHO-DXB11. In terms of copy number gains, CHO-S had the highest genes with copy number gain followed by CHO-DG44 and CHO-DXB11 where a major proportion of genes showed approximately a two-fold amplification across all three cell lines. Further, we mapped these genes to their differential expression profile which were categorized into following types: a) Concordant amplification/deletion - increase or decrease in gene expression due to concordant copy number gain or loss, respectively, b) Opposing amplification/deletion - increase or decrease in expression of the genes compensated by copy number loss or gain, respectively, and c) Silent amplification/deletion of gene copies (non-significant expression change with copy number change). Our result revealed that most of the genes belonged to the third category where average silent amplification was 88.66% ± 8.32% (mean ± SD) and average silent deletion was 87% ± 7% (mean ± SD) as the copy number gain or loss did not translate to the expected outcome and the expression profile of the corresponding genes remained unaffected.

As CHO-DG44 is known to have *Dhfr* gene deleted, we analysed the genes which showed copy number variations between CHO-K1 and -DG44 and noted that while *Dhfr* activity is found in several DG44 clones (due to reinsertion during selection process), the neighboring genes such as *Fam151b*, *Msh3* and *Zfyve16* have have been completely lost from the DG44 genome as reported previously (Lakshmanan et al. 2019). A similar result was also observed for CHO-DXB11 where only *Msh3* and *Zfyve16* showed copy number loss (Vishwanathan et al. 2017; Kaas et al. 2015). Interestingly, we also noted a significant concordant copy number loss shared between CHO-S and CHO-DG44, including genes such as *Chcr1*, *Mtss1* and *Mrpl13*. It should be noted that as these genes are not genomically clustered, i.e., from the same chromosome, it is highly unlikely that common genes have lost copies due to just by chance. It indicates that both CHO-DG44 and CHO-S could have emerged from a common ancestor, while still there is a possibility that convergent genomic instability or independent culture-adaptation selection could have also contributed to this shared loss.

Although CHO-S had more genes with copy number gain than CHO-DG44 and CHO-DXB11, most gains were not reflected in concordant increase in gene expression. Not only the concordant copy number gain and upregulation were more in CHO-DG44, but nearly 4.9% of the genes having copy number loss show increased differential expression, suggesting a compensatory mechanism. This indicates that while CHO-DG44 has more upregulated genes which are influenced by copy number gain, it also has a compensation response to copy number loss.

We also investigated the genes which showed silent amplification, i.e., genes with CNV gain but no significant gene expression change, where most of them had genetic information processing roles, e.g., *Gins2*, *Ddx24*, *Sbno2*, *Adat3*, *Foxo6*, *Mknk2*, and *Znf775*, or involved in secretory and vesicle trafficking pathways, along with cell adhesion and cell cycle regulation. Notably, CHO-S had most number of metabolic genes included in this category (*Acadvl*, *Aldh7a1*, *Bckdha*, *Cdo1*, *Idh3g*, *Ndufa2*, *Pcyox1l*, *Pgk1* and *Atp8b1*) indicating that metabolic activity is tightly regulated at transcriptional and post-transcriptional level despite genomic amplification. Together, these observations highlight that although copy number changes could explain the differential expression of a subset of genes, it represents only a smaller fraction of the entire dataset and is unlikely to be a major driver of the observed cell line heterogeneity.

### Analysis of CHO global promoter architecture reveals functional division between housekeeping and dynamically regulated genes

The absence of a clear relationship between CNV and gene expression indicates that the cell-line-specific expression divergence observed is more likely regulatory than genomic in origin. We therefore examined the differences in transcription initiation site, by specifically analysing the differential promoter usage. As a first step, we characterized the global promoter architecture of the CHO genome and identified 30,767 promoters were identified across 23,141 annotated genes using the CHO RefSeq annotations based on the assumption that transcripts which have identical or very close transcription start sites (TSSs) are regulated by the same promoter (see **Methods**). We then identified genes containing single versus multiple promoters and identified that a larger proportion of the genes, i.e 78.8 %, have only single promoters while 21.2% genes have multiple promoters (ranging from 2 to >5 promoters) (**Figure 5a, Supplementary file 5, Table 1-2**). We also estimated promoter activity as the total quantity of transcripts which are initiated at each promoter based on the set of unique junction reads from RNA-seq. Therefore, we found a perfect linear correlation between gene expression and activity of single promoters while a non-linear association was inferred in multi-promoters (**Figure 5b**)

**Fig 5.**
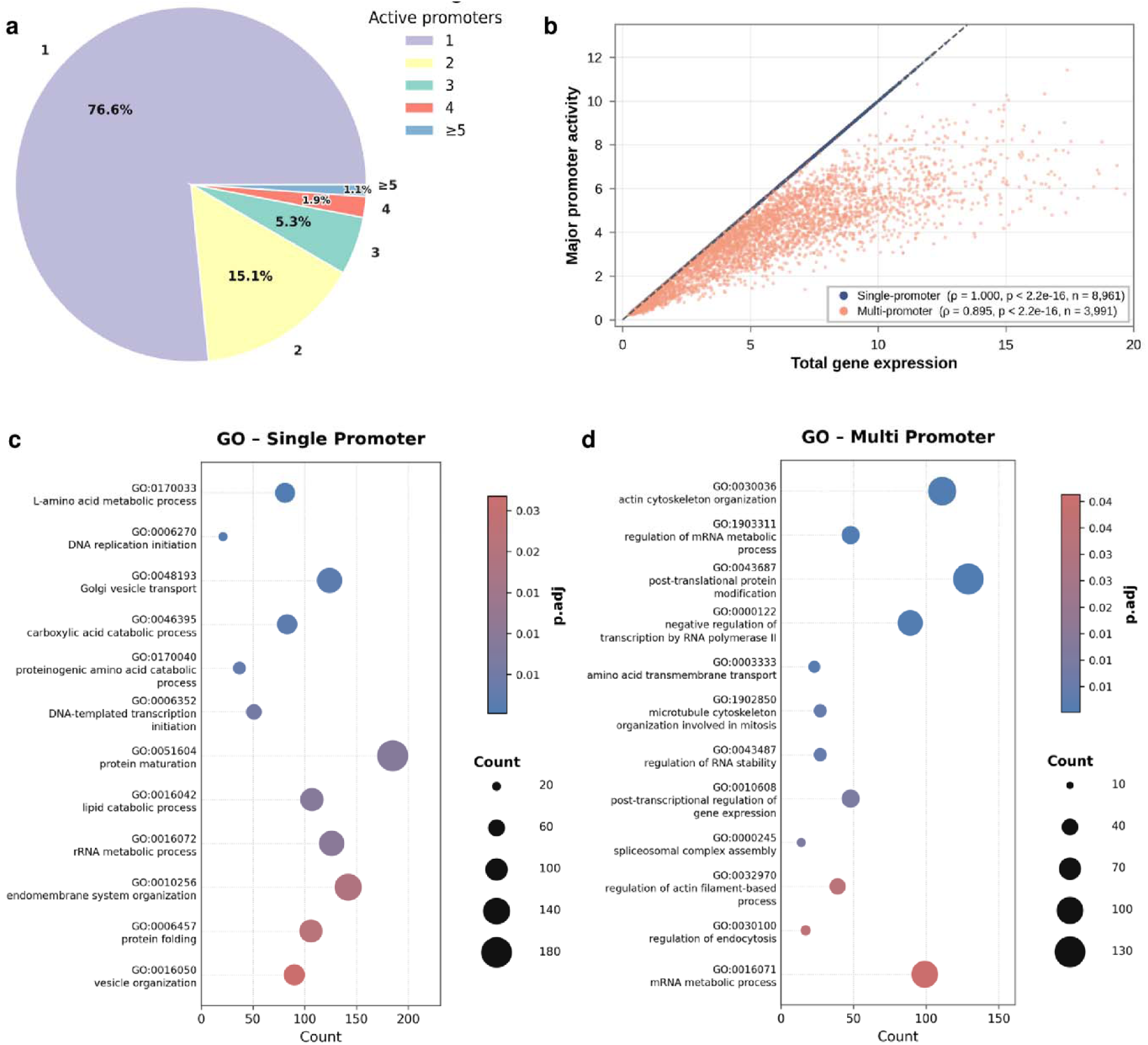
Promoter analysis reveals distinct functional roles for single and multi-promoter genes in CHO cell lines. (a) Proportion of genes by number of active promoters (1, 2, 3, 4, ≥5), as quantified using proActiv (b) Relationship between major promoter activity and total gene expression for single-promoter and multi-promoter genes (c,d) Functional enrichment analysis (GO/KEGG pathway) of gene categorized by promoter architecture. (c) Enrichment of pathways involved in housekeeping function among single-promoter genes. (d) Enrichment of pathways involved in context dependent control / require dynamic regulation among multi-promoter genes. P-values were adjusted using Benjamini - Hochberg correction and P-value < 0.05 was considered as statistically significant.

Enrichment analysis of single and multiple promoter genes revealed distinct functional biases. Genes having a single promoter have predominantly housekeeping functions and multiple metabolic pathways (**Figure 5c**). On the other hand, pathways controlled by multiple promoters were mostly involving epigenetic regulation (**Figure 5d**), consistent with previous studies which have also reported that alternative promoter usage (APU) is a key mechanism in controlling the expression of chromatin regulatory machinery of the cells (Hamaya et al. 2023). Similarly, actin cytoskeleton organization (GO:0030036; adj *P* < 0.01), post-translational protein modification (GO:0043687; adj *P* < 0.01), vesicle tethering complex (GO:0099023; adj *P* = 0.02) and regulation of endocytosis (GO:0030100; adj *P* = 0.03) pathways were also enriched in multi promoter genes **(Supplementary file 5, Table 3-4)**. The enrichment of these pathways together with the epigenetic remodeling related pathways in multi-promoter genes indicate that cellular processes which require dynamic and context-dependent control preferentially rely on alternative promoters to change the gene expression in a cell-type or condition-specific manner.

We also noted that pathways related to transcription are enriched in both single and multi-promoter genes where the functions of these pathways in each set vary greatly. For instance, genes involved in basal transcription factors (cge03022; adj *P* < 0.01) and RNA polymerase (cge03020; adj *P* = 0.02) are controlled by single promoters while post-transcriptional regulation of gene expression (GO:0010608; adj *P* < 0.01) and negative regulation of transcription by RNA polymerase II (GO:0000122; adj *P* < 0.01) are controlled by multi promoters. Besides that, genes related to splicing like RNA splicing (GO:0008380; adj *P* = 0.04) and spliceosomal complex assembly (GO:0000245; adj *P* = 0.01) also have multiple promoters. Consistent to this observation, it was previously reported that APU controls alternative splicing and generates different mRNA isoforms (Huin et al.,2017).

We next examined the promoter architecture of CHO housekeeping genes (Bahr et al. 2009; Ma et al. 2020), as it has been reported that their expression should be tightly regulated to facilitate stable expression; we hypothesized that these genes may employ single promoters to minimize variability in transcriptional output. Out of nearly 28 previously reported housekeeping genes, 22 were controlled by single promoters, coinciding with one of the previous reports which identifies the prevalence of single promoters (especially CpG rich) in housekeeping genes (Baek et al. 2007). Another interesting observation is that pathways related to viral infection were also significantly enriched among the genes having single promoters, indicating that similar to housekeeping functions, viral response genes may also be tightly controlled for their expression and reducing transcriptional noise.

### Promoter architecture and transcriptional initiation landscapes drives transcript isoform diversity across CHO lineages

Next, we quantified the activity of 30,767 promoters across different CHO cell lines by considering the amount of total transcriptional output initiated at each promoter using the set of unique junction reads as defined previously (Demircioğlu et al. 2019). We identified 9784 (31%) promoters as major promoters (most active promoters), while 12368 (40.2%) and 8615 (28%) promoters as minor and inactive promoters, respectively (**Figure 6a, Supplementary file 5, Table 5**). Then, we analysed the position of major promoters among multi-promoter genes and noted that while almost half of them had the canonical promoters (the first annotated promoter within the gene) as the major promoter (∼48%), we also found major promoters are located downstream of TSS as well (**Figure 6b**). Moreover, we also noted that as the number of promoters increases, the proportion of minor promoters increases at the expense of major and inactive promoters. This also indicates that the promoter choice is different for different genes and canonical promoters need not be the ones with the highest transcriptional activity.

**Fig 6.**
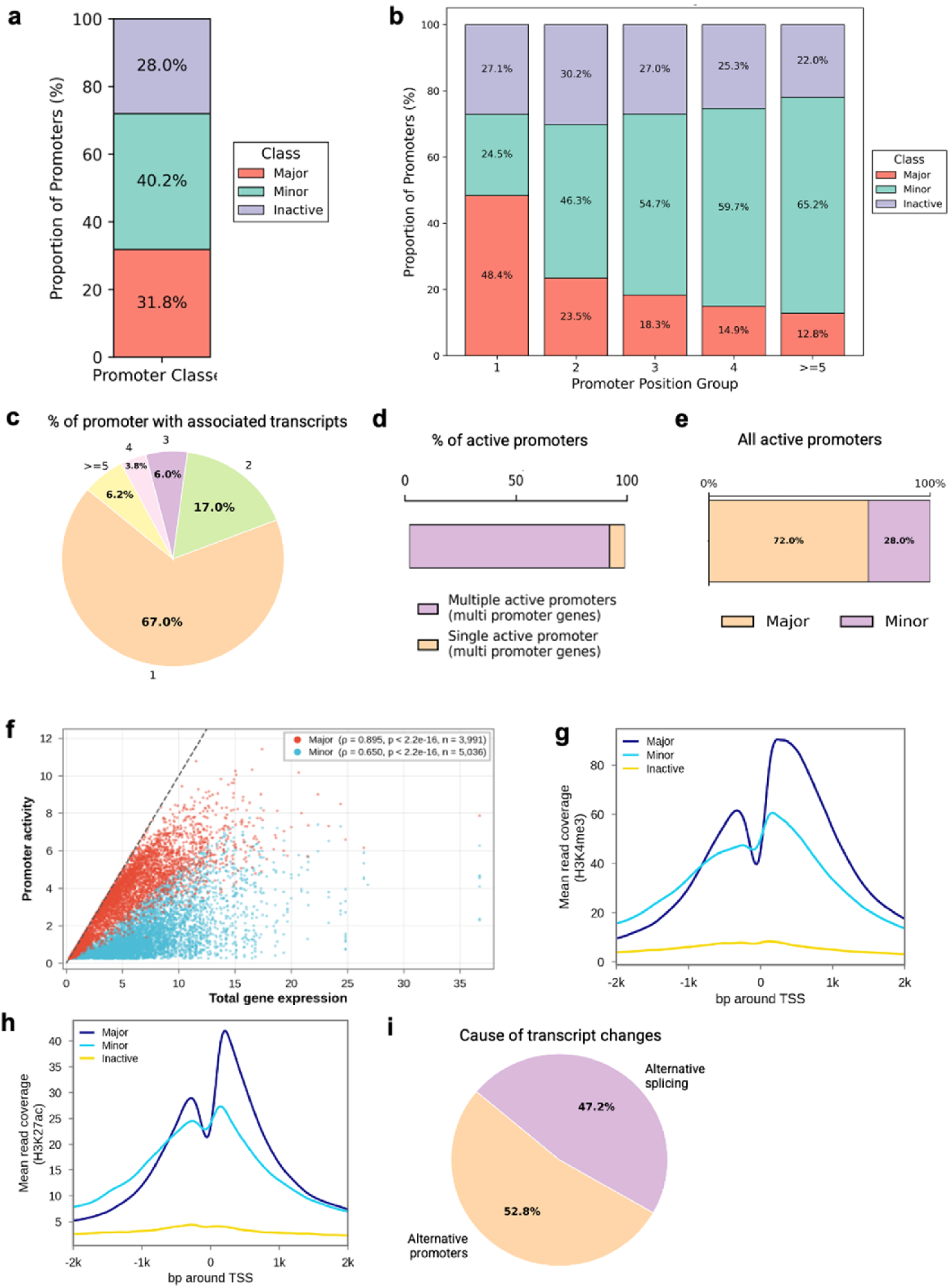
Promoter activity classes, transcript diversity and chromatin support in CHO cells. (a) Proportion of promoters classified as major, minor or inactive. (b) Proportion of major, minor and inactive promoters by promoter position within the gene (1, 2, 3, 4, ≥5). (c) Proportion of promoters associated with one, two, three, four or ≥5 transcripts. (d) Proportion of active promoters in multi-promoter genes that belong to genes with multiple active promoters versus genes with a single active promoter. (e) Proportion of major and minor promoters among all active promoters. (f) Promoter activity versus total gene expression for major and minor promoters. (g, h) Mean read coverage of H3K4me3 (g) and H3K27ac (h) ChIP-seq signal around the TSS for major, minor and inactive promoters in CHO-K1. (i) Cause of transcript differences in multi-promoter genes: alternative promoters or alternative splicing.

We then examined how active promoters are distributed among the expressed genes. Out of 12,952 genes with at least one active promoter, 3,329 (25.7%) used two or more active promoters, and 3,111 (24.0%) had at least two promoters each contributing more than 10% of the overall gene expression (**Figure 6c**). Among multi-promoter genes, 92.7% of the active promoters belonged to genes having more than one active promoter, while only 7.3% were the sole active promoter of their gene (**Figure 6d**). Across all 17,988 active promoters, 72.0% were major and 28.0% were minor promoters (**Figure 6e**), indicating that although most genes are driven by a single dominant promoter, a considerable set of genes employ additional promoters to initiate transcription. To assess how much each promoter class contributes to the transcriptional output, we compared the mean activity of major and minor promoters with the overall expression of their corresponding genes across all samples (**Figure 6f**). Major promoter activity was strongly correlated with gene expression (Spearman ρ = 0.895; n = 3,991), whereas minor promoter activity showed a weaker association (ρ = 0.650; n = 5,036). Although the major promoter provided at least half of the gene expression in 87.7% of multi-promoter genes, minor promoters collectively accounted for 34.2% of the total promoter activity within these genes. This indicates that for a substantial fraction of genes, gene-level quantification masks a considerable contribution from alternative promoters, which is not captured by standard differential gene expression analysis.

To further confirm the estimated promoter activities from RNA-seq data, we compared the promoter estimates to that of publicly available H3K4me3 and H3K27ac ChIP-seq data from the CHO-K1 cell line. This comparison indicated that the highest amount of mean H3K4me3 signal within 200 base pairs (bp) of the TSSs was observed for major promoters while inactive promoters showed the least H3K4me3 signals (**Figure 6g & 6h, Supplementary file 5, Table 5**). The differences among the three classes were significant for both marks (Kruskal–Wallis p < 2.2 × 10□¹□), including significant difference between major and minor promoters (Mann–Whitney p = 1.4 × 10□□² for H3K4me3 and 1.8 × 10□□□ for H3K27ac). These results confirm that RNA-seq based promoter activity estimates are quite robust as reported previously (Demircioğlu et al. 2019).

Since alternative promoters initiate transcription from distinct first exons, we next analysed their contribution to transcript isoform diversity in CHO cells. By mapping the RefSeq transcripts of multi-promoter genes to their respective promoters, we found that 67.0% of the promoters give rise to a single transcript, while the remaining 33.0% drive two or more transcript isoforms. To quantify the relative contribution of alternative promoter usage and alternative splicing, we compared all pairs of transcripts within each multi-promoter gene and attributed the difference to alternative promoters when the two transcripts were initiated from different promoters, and to alternative splicing when they shared the same promoter. Out of 72,012 transcript pairs, 38,022 (52.8%) differed by their promoter and 33,990 (47.2%) by splicing alone (**Figure 6i, Supplementary file 5, Table 6**), indicating that alternative promoters contribute slightly more than alternative splicing to transcript diversity in CHO cells. This is in line with the previous report in human tissues, where 58% of isoform switching events involved a change in promoter (Demircioğlu et al. 2019).

### Promoter activity analysis reveals lineage-specific transcriptional identity and buffering in CHO cells

Given that alternative promoters contribute significantly to transcriptional diversity in CHO cells, we investigated whether these promoter-level profiles reflect CHO lineage-specific gene expression differences. To first examine whether promoter activity captures the transcriptional identity of the CHO cell lines, we performed t-distributed stochastic neighbor embedding (t-SNE) using the activity of the promoters of all genes across all CHO samples. The t-SNE revealed a clear separation of samples according to their cell line, where CHO-DG44, CHO-S and CHO-DXB11 samples formed distinct clusters, whereas CHO-K1 samples were distributed across several clusters largely defined by their culture media (**Figure 7a**). This mirrors our observations from the gene expression analysis, indicating that cell line identity is a major source of variation in promoter usage across CHO cells, with culture media contributing additional variation, particularly among CHO-K1 samples. Hierarchical clustering of the mean activity of these promoters across the four cell lines revealed distinct groups of promoters that were specifically active in one cell line compared with the others (**Figure 7b**), indicating that each CHO cell line is characterized by a unique set of active promoters, even when the expression of some genes remains largely unchanged. We considered the contribution of single and multiple promoters separately in driving the cell line specific variation considering the promoter activity of CHO-K1 as reference. Out of 12545 unique promoters associated with 4916 genes that had multiple promoters, only the statistically significant ones without any internal promoters were considered for the analysis in respective cell lines. Further, promoter activity was categorized into two groups, with a log2FC ≥ 1 threshold defining upregulated activity and log2FC ≤ −1 threshold defining downregulated activity. In all cell lines, few proportions of multiple promoters showed contrasting activity, i.e., while one promoter showed upregulation others showed downregulation. We found 67 (3.23%), 64 (3.1%) and 43 (2.8%) such promoters, in CHO-D44, CHO-S and CHO-DXB11, respectively. After removal of these promoters showing contrasting activity, CHO-DG44 showed 786 (37.9%) and 796 (38.3%) promoters with up and downregulated activity, respectively. Similarly, CHO-S showed 609 (30%) and 928 (45.8%) up and downregulated promoter activity, respectively while CHO-DXB11 showed 401 (26.5%) and 817 (54.07%) promoters with up and downregulated activity, respectively (Supplementary file 5, Tables 7-9).

**Fig 7.**
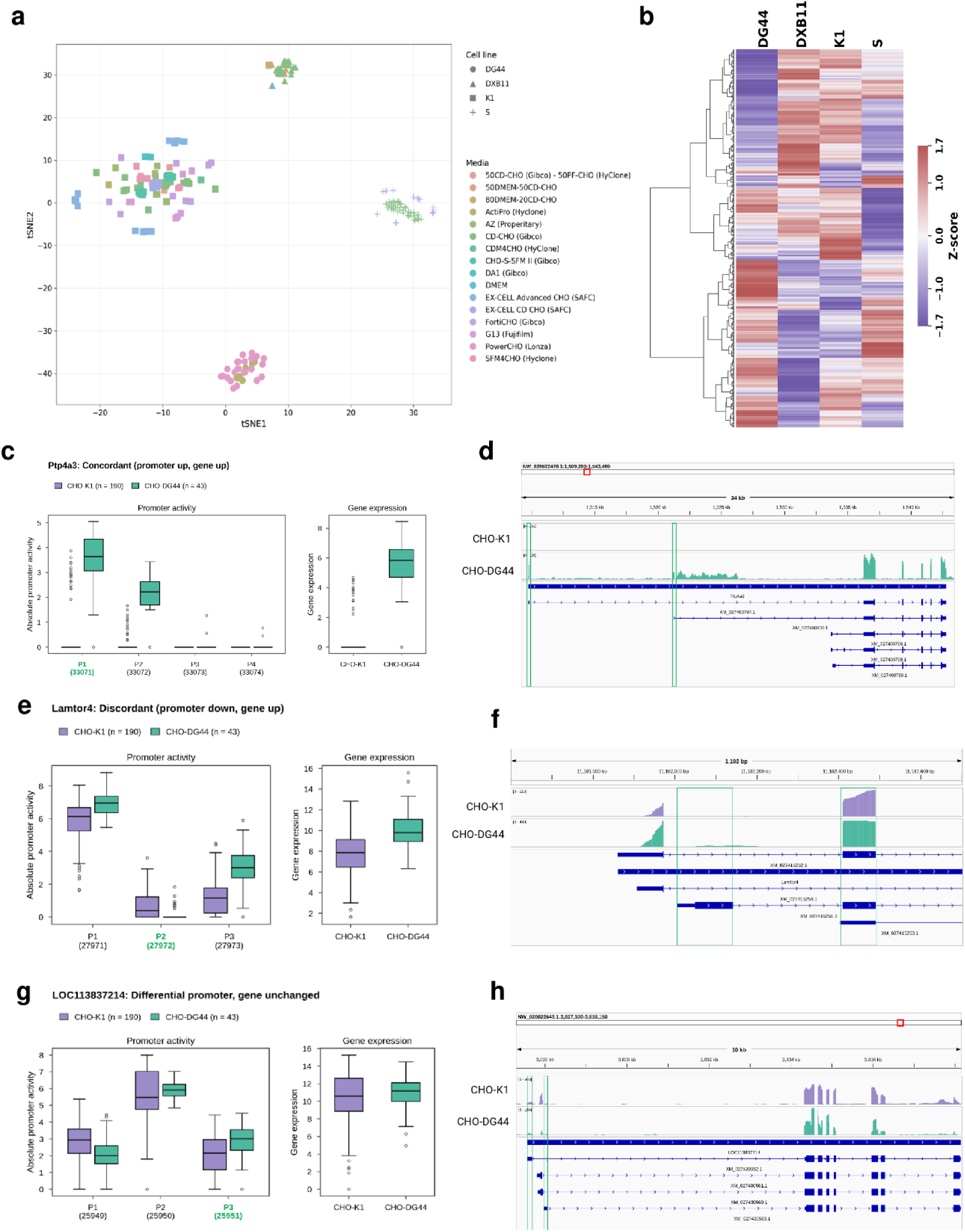
Cell line-specific promoter activity and its relationship to gene expression. (a) t-SNE of CHO samples based on promoter activity, coloured by medium and shaped by cell line. (b) Hierarchical clustering heatmap of the mean activity of cell line-specific promoters across CHO-DG44, CHO-DXB11, CHO-K1 and CHO-S. (c, e, g) Promoter activity (left) and gene expression (right) in CHO-K1 and CHO-DG44 for three example genes: *Ptp4a3* (concordant: promoter up, gene up), *Lamtor4* (discordant: promoter down, gene up) and *LOC113837214* (differential promoter activity, gene expression unchanged). (d, f, h) Read coverage tracks at the corresponding loci, with differentially active promoters highlighted.

Next, we shifted our focus to analyze the influence of promoter activity on gene expression by analyzing whether genes showing up/down regulation of promoter activity also show corresponding change in gene expression across the three cell lines and categorized into three types: 1) concordant relationship between promoter activity and gene expression, 2) discordant relationship between promoter activity and gene expression, and 3) promoter fold change without any influence of gene expression. Based on this analysis, we found that concordant events (both up and down regulation) were more as compared to discordant events. CHO-S had the highest number of concordantly upregulated genes (25.2%), while CHO-DG44, ranked second (21.4%) and CHO-DXB11 ranked third (10.11%) in this category. In terms of concordant downregulation CHO-DXB11 had the highest number of genes (17.55%). When compared to concordant events, discordant events (both up & down) were extremely low (0% to 0.5% only). Here, it should be noted that while the proportion of genes showing concordance between gene expressions and promoter activity is more than discordance (19-32% vs 0-0.5%), still the highest number of genes which had promoter switching did not contribute to gene expression change (57-67%). We have also mapped the promoters showing contrasting activity with their gene expression profile and observed that, for most of the genes, the overall gene expression appears to reflect the predominant direction of promoter activity, suggesting that the combined contribution of multiple promoters may determine the final expression outcome. For example, in -DG44 *Arhgef28*, *Col4a1* and *Arhgef10l* exhibited an overall decrease in gene expression consistent with the predominant downregulatory activity of its promoters. Altogether, evaluation of the promoter activity in CHO cell lines identified that gene expression in CHO cells are either reasonably well correlated with overall promoter activity although it is buffered by post-transcriptional or chromatin-level mechanisms.

## Discussion

CHO cells are widely used mammalian host cells for large scale production of biopharmaceuticals, however, a comprehensive understanding of their transcriptional behavior is still lacking. In the past, a handful of studies have previously assembled the transcriptome from multiple cell lines and media conditions but do not cover a wide range of conditions as well as do not compare the transcriptome of CHO cells with Hamster tissues. In the current study, we have compiled transcriptome data from multiple CHO cell lines grown in various conditions and media as well as 9 Chinese hamster tissues and an online CHO transcriptome compendium was built. To facilitate the broader utility of the datasets compiled, we have developed a web-based user interface.

By leveraging on the newly built CHO transcriptome compendium, we first analyzed the key traits of CHO cells when compared to the Chinese Hamster and how diverse the cell lines are. We noted that several viral receptors are either lost or reduced in CHO cell lines but not in tissues, indicating that the resistance to multiple viral infections is a CHO cell line specific trait than that of Chinese Hamsters in general. This may be an adaptive trait acquired due to their long-term *in vitro* culture in aseptic environments, conferring intrinsic resistance to viral contamination. We also noted a significant simplification in terms of glycosylation in CHO cells. The ability of CHO cells to perform human-like PTM, especially glycosylation, is one of the major reasons for widespread use of this cell line across biopharmaceutical industries. Though we did not find major changes between hamster and CHO cells in terms of N-glycosylation pathway gene expression, significant differences were found in O-glycosylation, especially GalNAc-type-O glycosylation. Consistent with previous reports, we also found reduced expression of several GalNAc-Ts involved in mucin-type O-glycan initiation and core extension (*Galnt* family, *C1galt1c1l*, *B3gnt3* and *Gcnt4*) in CHO cell lines while the expression is intact in hamster tissues (Ho et al. 2016; Xu et al. 2011; Yang et al. 2014; Riedl et al. 2025). The loss of gene expression of multiple independent parts of glycosylation including O-glycan initiation and core extension, glycosphingolipid biosynthesis and bisecting GlcNAc via *Mgat3* indicates that CHO cells could possess a narrower glycosylation repertoire than Chinese Hamsters. Notably, to the best of our knowledge, our study represents the first systematic comparison of potential glycosylation variation between CHO cells and Chinese Hamster tissues, although based on glycosylation-pathway related gene expression rather than at the glycan-structure level, further pointing towards the need for such comparative glycomics in the future. We also noted several endopeptidase inhibitors, mostly serine protease inhibitors, either lost expression or possess reduced expression in CHO cell lines. While the expression of these genes is largely restricted to hepatocytes and plasma, their reduced expression in ovary-derived CHO cells indicate the tissue origin specific loss which could have been further compounded by the immortalization event. Here, it should be highlighted that this finding is significant as it has been already well-documented that multiple CHO cathepsins B and D cause proteolytic fragmentation and quality defects in recombinant antibodies during cell culture and downstream processing (L. Hu et al. 2022). Reduced endogenous serpin-mediated inhibition as well as removal of serum, a natural protease inhibitor in cell culture, leave the recombinant proteins produced from CHO cells completely unproductive, and thus the addition of endopeptidase inhibitors could increase the quality of recombinant proteins. Moreover, future studies could focus on profiling the simultaneous protease activity and endogenous serpin inhibitor expression levels to unravel this cross talk more clearly.

The contribution of cell line lineage to the CHO transcriptome is well documented, and our findings are also in line with previous observations where CHO-K1 and CHO-DXB11 exhibit similar global gene expression patterns while CHO-S and -DG44 showed distinct gene expression profiles. Such lineage-specific transcriptional signatures have direct implications in cell line selection for bioprocess development. For instance, the upregulation of several cadherin and claudin genes in CHO-DG44 highlights its high propensity for cell aggregation/clumping although it is the general tendency of cells to clump when grown in suspension cultures. While cellular clumping is a recognized problem in CHO culture, studies on this topic comparing different cell lines or media is relatively scarce; the only available report described DG44 cells were less aggregation-prone than CHO-K1 under standard conditions, while TNF-α induction increased aggregation in DG44 more than in K1 (Louie et al. 2020). As all samples analysed here are from serum-free cultures and media composition itself is known to affect cellular aggregation (Green et al. 2025; Boraston et al. 1992), media design could be an important lever for managing this tendency in CHO-DG44. We also noted the higher sensitivity of CHO-S and CHO-DG44 to amino acid deprivation which point towards more deliberate cell-line specific feeding strategies and media formulations to sustain high viability and productivity over extended culture periods. In addition to cell lines, we also found cell culture media to have a significant contribution to global transcriptome variation in CHO cells. A few previous studies have evaluated the performance of different media and have benchmarked them based on cell growth and mAb productivity (Rodrigues et al. 2012; Reinhart et al. 2015), while the underlying transcriptional responses remain unexplored. In this regard, although our compendium pointed towards broader media-induced effects, the analysis is inherently limited by the heterogeneous nature of datasets, data is acquired from different cultivation modes, scales and more importantly the media were adapted and optimized to the clones only in some instances (Sedighikamal et al. 2022; Rodrigues et al. 2012; Gyorgypal et al. 2025; Dhulipala et al. 2011). Therefore, to study the exact effect of media, a tightly controlled analysis needs to be done where cell lines will systematically be grown in different media compositions.

Establishing a direct relationship between genomic variation and transcriptomic profile has always been a major challenge. Although genomic architecture provides a blueprint of transcriptional output, gene expression is controlled by complex interplay among several factors such as epigenome context, and pre-and post-transcriptional modifications. In this complex landscape, the impact of CNVs is considered as an important genomic contributor though its precise role remains debated. While a few studies have earlier reported CNV to directly affect gene expression (Shao et al. 2019), few others have also pointed out the non-linearity of relationship between these two (Goswami et al. 2026; Ronemus et al. 2025). A handful of studies have earlier investigated the contribution of gene copy number to transcriptional output in CHO cell lines (Lakshmanan et al. 2019; Vishwanathan et al. 2017; Lewis et al. 2013; Huhn et al. 2022), yet whether CNV actually drives the transcriptional divergence observed across CHO cell lines remains unresolved. We therefore studied the role of CNVs in driving cell line specificity and found that a larger proportion of CHO genes either do not show any copy number alteration or their expression do not get affected by the CNVs. While a subset of genes whose expression was concordant with with copy number gain/loss, another small subset of genes, mostly related to epigenetic regulation, showed discordant relationship, indicating potential gene dosage compensation as seen in several cancer cells primarily in response to aneuploidy owing to their high genomic instability (Bravo-Estupiñan et al. 2023). As CHO cells are also characterized by prominent aneuploidy and genomic instability (Vishwanathan et al. 2017; Deaven and Petersen 1973; Yamano-Adachi et al. 2023), a similar dosage-compensation mechanism may underlie the discordant events observed here. Collectively, our analysis indicates that CNV contributes minimally to the transcriptional variation observed across CHO parental cell lines, highlighting the alternative regulatory mechanisms as primary drivers.

We also analyzed the role of promoter activity in driving cell line specific variance and noted a few key observations. First, we noted the promoter architecture in CHO cells is functionally stratified where the single-promoter genes belong to housekeeping and core metabolic functions, while multi-promoter genes were enriched for epigenetic, cytoskeletal and trafficking functions that require context-dependent control. Importantly, we noted ∼80% of CHO housekeeping genes (22 of about 28) were single-promoter genes, consistent with previous reports that constitutively expressed genes favor simple CpG-rich promoter architectures (Baek et al. 2007). Overall, a large number of genes (78%) were driven by single promoter whose expression scaled linearly with the promoter activity, while only a smaller proportion of genes (∼22%) were driven by multiple promoters though this could be a conservative estimate, as it depends on RefSeq annotation depth for the Chinese hamster available in NCBI. Further analysis of multiple promoter genes indicated that less than half of them (48%) were driven by the first canonical promoter as a major promoter, while the minor promoters accounted for 34% of the total expressed multi-promoter genes. Alternative promoters also accounted for slightly more transcript diversity than splicing alone (52.8% vs 47.2% of transcript pairs), comparable to the 58% reported for human tissues (Demircioğlu et al. 2019). These observations indicate that promoter choice may therefore be an underappreciated source of isoform variation in CHO, with potential consequences for N-terminal protein variants and regulatory UTR content. Finally, we noted that promoter activity was able to separate the cell lines as it was in gene expression, indicating that promoting switching events are a potential contributor to observed gene expression variations in CHO cell lines. While concordant promoter activity-gene expression changes greatly outnumbered discordant ones (19-32% vs 0-0.5%), more than half (57-67%) of differentially active promoters did not change gene expression. Importantly, we also noted that where promoters within a gene moved in opposite directions, overall expression also generally followed the dominant direction although the results are not statistically significant, which could be because of the different cut-offs that we used for such analysis. Overall, the promoter analyses indicated that even though promoter switching could be a potential contributor to gene expression divergence observed among cell lines, it is frequently buffered at the post-transcriptional or perhaps at chromatin level itself. Moreover, there could be other potential contributions from epigenetic regulation, splicing, transcription factor activity and mRNA stability which were not analysed in this study.

In conclusion, the current work bridges a critical knowledge gap between native Chinese hamster physiology and industrially relevant CHO cell lines using the transcriptome compendium. Our work particularly demonstrates how alternative promoter architecture and post-transcriptional buffering, rather than simple gene dosage variations, dominate in shaping cellular phenotypes. This suggests that the phenotypic variability observed across CHO lineages is mostly driven by dynamic regulatory rewiring while retaining core cell functionalities. Overall, we believe that the extensive dataset and web resource presented in this study will act as a launchpad for CHO synthetic biology and precision cell engineering initiatives, paving the way for the next-generation rational design of high-performance biomanufacturing platforms.

## Materials and Methods

### RNA-seq data collection and curation for compendium development

RNA-seq datasets were obtained from the Sequence Read Archive (SRA) and the European Nucleotide Archive (ENA). Studies were included if they profiled primary Chinese hamster tissue or one of the four parental CHO cell lines (CHO-K1, CHO-S, CHO-DG44 or CHO-DXB11), regardless of culture conditions, growth phase or recombinant protein expression status. In total, 23 unique BioProjects from 24 studies were selected, comprising 350 RNA-seq samples. In addition to this publicly available data, unpublished in-house RNA-seq of six Chinese hamster tissues (brain, kidney, liver, lung, ovary and spleen) were also included together with public data. Together, RNA-seq of nine Chinese hamster tissues, brain, heart, kidney, liver, lung, muscle, ovary, pancreas and spleen, are included in this collection. CHO samples represented varying bioprocessing conditions, ranging from adherent and suspension cultures, batch and fed-batch processes, serum-containing and serum-free media, and parental or recombinant-protein-producing (producer vs. non-producer) cells. Note that CHO-K1 samples include both the CHO-K1 and CHO-K1-GS cell lines as well. Information regarding accession numbers of all samples together with their library layout, strandedness, cell culture medium, cell line, producer status and corresponding journal publications (if any), are listed in the sample metadata table (**Supplementary file 1, table 1**).

### RNA-seq processing and quantification

Read quality was assessed with FastQC v0.11.9 (Andrews 2010) before and after trimming. Adapters and low-quality bases were removed with Trimmomatic v0.39 (Bolger et al. 2014) using ILLUMINACLIP. The compendium mixes single-end and paired-end libraries as well as poly(A)-selected, total-RNA and rRNA-depleted protocols, so adapter sequences differed between studies. Where non-standard adapters were present, Cutadapt v3.5 (Martin 2011) was applied. Samples with abnormal quality profiles were inspected manually before inclusion. Trimmed reads were then aligned to the Chinese hamster genome CriGri-PICR (GCF_003668045.1) (Rupp et al. 2018) with HISAT2 v2.2.1 (Kim et al. 2019). The genome index was built with the splice-site and exon information from the matching RefSeq annotation (GCF_003668045.1_CriGri-PICR_genomic.gtf). Library strandedness was set from the metadata of the original studies. Reads were assigned to genes with featureCounts (Subread v2.0.3 (Liao et al. 2014), counting uniquely mapped reads against the CriGri-PICR annotation, with settings appropriate to single-end or paired-end libraries.

### Normalisation and global expression analysis

Raw RNA-seq counts were normalised with edgeR (Robinson et al. 2010) using the trimmed mean of M-values (TMM) (Robinson and Oshlack 2010). For this purpose, a multifactor design matrix incorporating cell line, cell culture medium, cell producer status (recombinant producer vs. non-producer), batch, sequencing layout, and library was used to estimate common dispersion, and the resulting normalized counts were used for further analysis. The normalized counts were then transformed as log2(counts + 1) and used for clustering and visualisation, expression summaries and the comparison of CHO cells with Chinese hamster tissues. Multidimensional scaling was performed using Pearson-correlation distances and non-metric isoMDS implemented in magrittr package in R, and the first two dimensions are plotted. For CHO samples, we visualized the normalised log2(counts + 1) values by UMAP (McInnes et al. 2018) using the uwot package in R with default settings. The cumulative transcriptional output was calculated as the fraction of total counts as a function of log2(counts + 1) for each cell type and tissue. Genes with near-zero expression in all samples were identified as genes with at most one read in any sample (n = 357) and tested for GO enrichment with clusterProfiler. To identify the median gene expression across tissues and cell lines, we followed the following procedure. A gene was considered expressed in a condition when its normalised log2 counts per million (log2CPM) was above 0 (about 1 CPM) in more than 95% of the samples of that condition. For each sample, the median of log2(TMM-CPM + 1) was computed, using TMM factors calculated on the raw counts and effective library size = library size × TMM factor, and compared between tissue and CHO samples with a two-sided Mann-Whitney U test.

### Variance partitioning analysis

To determine the contribution of experimental variations to gene expression changes across CHO samples was estimated with a linear mixed-effect model using variancePartition R package (Hoffman and Schadt 2016). The batch effect was fixed while phase, cell line, producer status, culture medium, library, and sequencing layout was modeled as random effects to quantify their contributions to gene expression variability. The analysis covered genes that have expression exceeding CPM>1 in at least 1% of samples (n = 20,774). Subsequently, genes with at least 95% of variance explained by cell line or media alone were highlighted as cell-line-specific and media-specific examples, respectively. Genes in which cell line and culture medium together explained a cumulative variance of at least 70% were taken forward for enrichment analysis.

### Differential expression analysis

#### CHO cells versus Chinese hamster tissues comparisons

In the compiled dataset since the number of CHO cell line samples was very high than Chinese hamster tissue samples, the CHO vs. tissue comparison was run in four independent sampled subsets. We specifically sampled 16 CHO samples four times and compared them with the same 16 tissue samples: subset 1, CHO-S; subset 2, CHO-K1 and CHO-S; subset 3, CHO-K1 and CHO-DXB11; subset 4, CHO-DG44. From each of the comparisons, genes were assigned into categories based on their expressions in the two groups following the RNA-based classification of genes used for the human tissue transcriptome (Uhlén et al. 2015). A gene was categorized as expressed in a group when its log2 value was above 1 in at least 85% of samples and as non-expressed when it was 1 or below in at least 85% of samples. Genes expressed in tissues and non-expressed in CHO cells were classified as completely lost in CHO, and genes non-expressed in tissues and expressed in CHO cells were classified as completely gained in CHO. Genes whose expression were reduced in CHO were identified by routine differential expression analysis with limma (Ritchie et al. 2015), a linear model with a CHO versus tissue contrast was fitted, moderated t-statistics were computed with eBayes, and genes with log2 fold change below −1 and a Benjamini-Hochberg adjusted P value below 0.05 were called reduced. The final gained, lost and reduced gene lists are the genes assigned to the same category in all four subsets. For the gained and lost gene sets, expression in tissues and CHO samples was compared per subset with two-sided Mann-Whitney U tests (16 CHO versus 16 tissue samples) on log2(normalised counts + 1). P-values were Holm-adjusted across all gene-by-subset comparisons within a figure, and the significance stars shown are those of the least significant subset for each gene. ***Expression differences between CHO cell lines:*** Cell-line comparisons used DESeq2 (Love et al. 2014) on raw counts. Note that CHO-K1 and CHO-K1 (GS) samples were merged into a single CHO-K1 reference group. Samples of CHO-K1, CHO-S, CHO-DXB11 and CHO-DG44 were analysed in one model with design ∼ cell line and CHO-K1 as the reference level. Genes with counts<10 in total were removed. The Wald test was applied to the contrasts CHO-S, CHO-DXB11 and CHO-DG44 versus CHO-K1, and genes with a Benjamini-Hochberg adjusted P value below 0.01 and an absolute log2 fold change above 2 were called differentially expressed.

#### Expression differences in CHO-K1 based on cell culture medium

The effect of culture medium was assessed within CHO-K1 samples only and the design ∼ medium. Nine media (ActiPro, CDM4CHO, CHO-S-SFM II, DA1, EX-CELL Advanced CHO, EX-CELL CD CHO, FortiCHO, G13 and SFM4CHO) were compared where each of them against CD-CHO (Gibco), which was chosen as the reference because it is a long-established chemically defined formulation. Genes with counts<10 in total were removed, Cook’s outlier refitting was enabled, and genes with adjusted P below 0.05 and an absolute log2 fold change above 1 were called differentially expressed.

### Functional enrichment analysis

Enrichment of Gene Ontology (biological process, molecular function, cellular component) and KEGG pathways were done using clusterProfiler v4.10.0 (Wu et al. 2021). For KEGG pathways, gene symbols were converted to Entrez identifiers and the enrichKEGG package was implemented with organism code “cge”. Terms were called significant if Benjamini-Hochberg adjusted P-value is below 0.05 (and q-below 0.05 for GO). A rich-factor cutoff of at least 10% (the fraction of a pathway’s genes present in the input set) was applied as the standard. For the tissue versus CHO comparison, a relaxed cutoff below 10% was adopted so that biologically important pathways that would otherwise be excluded by stringent filtering were retained. The same relaxed cutoff was used for the promoter-specific analyses, to allow a more inclusive assessment and capture potentially relevant changes. GO terms for *Cricetulus griseus* were taken from the AnnotationHub OrgDb record AH120126 (Shepherd, L, Morgan, M 2025) with gene symbols as the key type.

### Metabolic and secretory pathway mapping

Differentially expressed genes were mapped to the genome-scale metabolic network *i*CHO3K (Di Giusto et al. 2026), which resulted in 2,582 mapped genes, and the numbers of up-and down-regulated metabolic genes were counted per cell line. Lost and reduced genes from CHO-versus-tissue comparison were mapped to SecRecon, a curated reconstruction of the mammalian secretory pathway (Masson et al. 2025), at the system and subsystem levels, with subsystem names refined for specificity (for example N-glycosylation and O-glycosylation instead of glycosylation as it is in SecRecon). Over-representation of each system and subsystem was tested with a one-sided hypergeometric test (equivalent to a one-sided Fisher exact test), with the universe defined as the 965 unique CHO genes in the SecRecon mapping. For each term, *n* was the number of query genes in the universe, *k* the number of those in the term, *K* the number of all genes in the term and *M* the universe size; the enrichment ratio was (*k*/*n*)/(*K*/*M*) and the P value was *P*(*X*≥*k*). Benjamini-Hochberg adjusted values were computed within each table.

### Gene copy number variation

Gene copy number changes between the CHO cell lines were estimated calculated from normalized read depth of each gene using CHO-K1 as reference. Median normalized read depths were obtained for all genes across all cell lines from a previous study. An average value of CHO-K1 PF, CHO-K1 SF and CHO-K1 ATCC was used to estimate the gene read depth in CHO-K1 and the ratio of normalized read depths between CHO-K1 and other CHO cell lines, i.e., CHO-S, CHO-DXB11, CHO-DG44, was calculated. Genes were labelled as normal for ratios from 0.5 to 2. Gains were assigned to amplification classes of 2x, 4x, 6x and 8x by rounding the ratio down to the nearest even multiple, with 8x as the highest class. Losses were assigned using the inverse ratio, with classes 0.5x, 0.25x and 0.125x for successive doublings, and 0.0625x for an inverse ratio of 16 or more.

Gene copy number calls were then matched by gene symbol to the DESeq2 results for each cell line (CHO-S, CHO-DXB11 and CHO-DG44) compared with CHO-K1.The expression change was called up-regulated or down-regulated at adjusted P-value below 0.01 and absolute log2 fold change above 2. Each gene was then assigned to one of three categories: (a) concordant, where copy number gain coincides with up-regulation or copy number loss with down-regulation; (b) discordant, where copy number and expression change in opposing directions (amplified but down-regulated, or deleted but up-regulated); and (c) silent, where gene expression is not affected by the copy number change.

### Promoter activity analysis

#### Identification of promoters from genome annotation

Promoters were defined from the CriGri-PICR RefSeq annotation with preparePromoterAnnotation function in proActiv tool v1.6.0 (Demircioğlu et al. 2019), using the *Cricetulus griseus* species setting. Transcripts with identical or overlapping first exons were grouped under one promoter, and the 5’ end of the most upstream first exon was taken as the transcription start site. This identified 30,767 promoters associated with 23,141 genes. Running proActiv on this genome required a custom GenomeInfoDb file built from the CHO gene annotation and a CHO-specific BSgenome package (BSgenome.Cgriseus.NCBI.CriGriPICR) created with BSgenomeForge. The annotation object was built once and used for all samples. Within each gene, promoters were numbered from the 5′ end by proActiv (promoter 1 is the most upstream), and promoters at position five or beyond were grouped as “5 or more”. Genes were classified as single-promoter or multi-promoter according to the number of annotated promoters.

#### Estimation of promoter activity

proActiv was run separately on each HISAT2 BAM file. proActiv models each gene as a splicing graph, and the count of a promoter is the number of reads spanning the first junction (first intron) of the transcripts that use that promoter. Promoters with missing counts in a sample were removed, and the per-sample tables were merged by promoter identifier into one table of raw promoter counts. Absolute promoter activity was calculated for each sample by dividing the raw promoter count by the library size of that sample (the sum of all its promoter counts, in millions), which gives counts per million (CPM), and transforming the result as log2(CPM + 1). Gene-level counts were obtained by summing the raw counts of all promoters of a gene and were normalized in the same way, separately from the promoter counts.

#### Classification of promoters as major, minor or inactive promoters

For each gene, the promoter with the highest mean absolute activity across samples was called the major promoter, while promoters with mean absolute activity below 0.25 were called inactive, and the remaining promoters minor. The relationships between major or minor promoter activity and total gene expression were summarised with Spearman correlation. For multi-promoter genes, the proportions of major, minor and inactive promoters were tabulated by promoter position (1, 2, 3, 4 and 5 or more), and the number of transcripts per promoter was counted.

#### ChIP-seq analysis to validate promoter class predictions

Promoter classes identified by proActiv analysis were compared with publicly available H3K4me3 and H3K27ac ChIP-seq data from CHO-K1 cells grown in batch culture (ENA project PRJEB9291; time points T5, T9, T13 and T17) (Hernandez et al. 2019). Signal tracks (MACS2 treatment pileup) on the CriGri-PICRH assembly were scaled by the total signal of each track to correct for sequencing depth and averaged over the four time points. Because the ChIP-seq data are on CriGri-PICRH and the promoters were defined on CriGri-PICR, promoter positions were converted between the two assemblies using RefSeq transcripts present in both annotations, with local offsets taken from the nearest promoters that have a matching transcript. For each promoter, the mean signal within 1 kb on either side of the transcription start site was calculated; promoters on scaffolds without matching transcripts (about 2.5%) could not be placed and were left blank. Signal was compared among major, minor and inactive promoters with a Kruskal–Wallis test, and major and minor promoters were compared with a two-sided Mann–Whitney U test (**Supplementary file 5, Table 5**).

#### Contribution of promoters and splicing to isoform diversity

To determine the contribution of promoters and splicing to isoform diversity, first, RefSeq transcripts of multi-promoter genes were assigned to their promoters. All transcript pairs within a gene were then compared: a pair was assigned to alternative promoter usage if the two transcripts come from different promoters, and to alternative splicing if they share the same promoter.

#### Estimation of differential promoter activity

Differential promoter activity was estimated using DESeq2 (Love et al. 2014) on raw proActiv promoter counts using CHO-K1 as reference. Gene-level counts were the sum of the counts of all promoters of a gene. Features were kept if they had at least 10 counts in at least 20 samples, which left 14,305 of 30,767 promoters and 11,270 of 23,141 genes. A negative binomial model with design ∼ cell line was fitted separately at the promoter and gene levels, and the contrasts CHO-S, CHO-DG44 and CHO-DXB11 versus CHO-K1 were tested with the Wald test. Features with adjusted P-values below 0.05 and absolute log2 fold change above 1 were called significant. Cell lines were compared by t-SNE (Rtsne; perplexity 30, 1,000 iterations) on the log2(CPM + 1) activity of all promoters, and cell-line-specific promoters were shown as a heatmap of log2(CPM + 1) promoter activity, z-scored per promoter and clustered hierarchically (Euclidean distance, average linkage).

#### Alternative promoter analysis

The alternative promoter analysis was restricted to genes with more than one annotated promoter (12,542 promoters in 4,916 genes). Promoters were first filtered on statistical significance and internal promoters were then removed. Promoter activity was classified from the log2 fold change relative to CHO-K1: values above 1 were called up-regulated, values below −1 down-regulated, and values between −1 and 1 were considered to show no substantial change. For cell-line-specific promoter activity, promoters of multi-promoter genes that showed contrasting activity, with both an up-regulated and a down-regulated promoter in the same gene, were removed before the promoter changes were mapped to gene expression changes in each cell line separately. Percentages are relative to the number of significant promoters for that cell line.

#### Concordance between promoter and gene expression

Differential promoter activity was mapped to the gene-level calls from the same DESeq2 analysis for each cell line comparison with CHO-K1 separately. A promoter and gene change were called concordant when both had a log2 fold change above 1 (concordant up-regulation) or both below −1 (concordant down-regulation). They were called discordant when promoter activity had a log2 fold change above 1 and gene expression below −1 (discordant promoter up-regulation), or the reverse. Promoters showing differential activity without an effect on gene expression were called “promoter fold change without any influence of gene expression”. For the example genes (Ptp4a3, Lamtor4 and LOC113837214), promoter activity and gene expression were shown as boxplots of log2(CPM + 1), and coverage tracks combined RNA-seq signal with H3K4me3 and H3K27ac ChIP-seq signal from CHO-K1

## Code and data availability

- The raw sequencing data analysed in this study are publicly available from SRA and ENA under the study accessions listed in Supplementary file 1, Table 1.
- The compendium can be explored through an interactive web application at https://huggingface.co/spaces/SBCE-Lab/CHO-Web-App.
- All analysis scripts are available at https://github.com/SBCELab-IITM/CHO-transcriptome-compendium.
- Any additional information required to reproduce this work is available from the lead contact.

## Acknowledgements

We thank the Centre for Integrative Biology and Systems medicinE (IBSE) and Wadhwani School of Data Science and Artificial Intelligence (WSAI), Indian Institute of Technology Madras, for their support and for providing the computational resources used in this work. M.L. acknowledges the funding support from the Indian Institute of Technology Madras for funding support through the New Faculty Scheme (IP23242264BTNFSC009058) and Anusandhan National Research Foundation through Prime Minister Early Career Research Grant (ANRF/ECRG/2024/002538/LS). D.M. and M.A.S.P.M. acknowledge the Half-Time Research Assistantship (HTRA) from the Ministry of Education, Government of India.

## Author contributions

Conceptualization - M.L., D.M., D.S.; Methodology - D.M., D.S., M.L., M.M.; Software - D.S., P.V.V., N.G., R.R.; Formal Analysis, D.M., D.E.K., M.A.S.P.M.; Investigation, D.M., D.S., J.K., K.T.P.; Data Curation, D.M., D.S., M.M., M.L.; Writing – Original Draft - D.M., D.S.; Writing – Review & Editing - M.L., J.K., K.T.P.; Funding Acquisition - M.L., Supervision - M.L., J.K., K.T.P.

## Declaration of generative AI and AI-assisted technologies in the writing process

During the preparation of this work, the authors used Claude and ChatGPT in a limited manner to improve the language and readability of the manuscript. After using this tool, the authors reviewed and edited the content, and is fully responsible for the final content in this article.

## Declaration of Competing Interests

The authors declare no competing interests.

## Notes

https://huggingface.co/spaces/SBCE-Lab/CHO-Web-App

### Competing Interest Statement

The authors have declared no competing interest.

## References

Andrews, S. 2010. “FastQC: A Quality Control Tool for High Throughput Sequence Data.” https://www.bioinformatics.babraham.ac.uk/projects/fastqc/.

Baek, Daehyun, Colleen Davis, Brent Ewing, David Gordon, and Phil Green. 2007. “Characterization and Predictive Discovery of Evolutionarily Conserved Mammalian Alternative Promoters.” Genome Research 17 (2): 145–155.

Bahr, Scott M., Trissa Borgschulte, Kevin J. Kayser, and Nan Lin. 2009. “Using Microarray Technology to Select Housekeeping Genes in Chinese Hamster Ovary Cells.” Biotechnology and Bioengineering 104 (5): 1041–1046.

Berting, Andreas, Maria R. Farcet, and Thomas R. Kreil. 2010. “Virus Susceptibility of Chinese Hamster Ovary (CHO) Cells and Detection of Viral Contaminations by Adventitious Agent Testing.” Biotechnology and Bioengineering 106 (4): 598–607.

Bolger, Anthony M., Marc Lohse, and Bjoern Usadel. 2014. “Trimmomatic: A Flexible Trimmer for Illumina Sequence Data.” Bioinformatics (Oxford, England) 30 (15): 2114–2120.

Bravo-Estupiñan, Diana M., Karol Aguilar-Guerrero, Steve Quirós, et al. 2023. “Gene Dosage Compensation: Origins, Criteria to Identify Compensated Genes, and Mechanisms Including Sensor Loops as an Emerging Systems-Level Property in Cancer.” Cancer Medicine 12 (24): 22130–22155.

Chantranupong, Lynne, Rachel L. Wolfson, Jose M. Orozco, et al. 2014. “The Sestrins Interact with GATOR2 to Negatively Regulate the Amino-Acid-Sensing Pathway Upstream of mTORC1.” Cell Reports 9 (1): 1–8.

Cherian, Reeja Maria, Chunsheng Jin, Jining Liu, Niclas G. Karlsson, and Jan Holgersson. 2015. “A Panel of Recombinant Mucins Carrying a Repertoire of Sialylated O-Glycans Based on Different Core Chains for Studies of Glycan Binding Proteins.” Biomolecules 5 (3): 1810–1831.

Chiang, Austin W. T., Shangzhong Li, Benjamin P. Kellman, et al. 2019. “Combating Viral Contaminants in CHO Cells by Engineering Innate Immunity.” Scientific Reports 9 (1): 8827.

Deaven, L. L., and D. F. Petersen. 1973. “The Chromosomes of CHO, an Aneuploid Chinese Hamster Cell Line: G-Band, C-Band, and Autoradiographic Analyses.” Chromosoma 41 (2): 129–144.

Demircioğlu, Deniz, Engin Cukuroglu, Martin Kindermans, et al. 2019. “A Pan-Cancer Transcriptome Analysis Reveals Pervasive Regulation through Alternative Promoters.” Cell 178 (6): 1465–1477.e17.

Dhulipala, Prasad, Margaret Liu, Shannon Beatty, et al. 2011. “Differential Cell Culture Media for Single-Cell Cloning Requirements of CHO-S and CHO DG44 Cells.” Preprint, December. https://eu-assets.contentstack.com/v3/assets/blt0a48a1f3edca9eb0/bltcb7f58ef4aa49669/658c32938df436040adde4eb/BPI_A_120911AR11_O_170635a.pdf.

Di Giusto, Pablo, Dong-Hyuk Choi, Athanasios Antonakoudis, et al. 2026. “A Community Reconstruction of Chinese Hamster Metabolism and Structural Systems Biology Elucidate Metabolic Rewiring in Lactate-Free CHO Cells.” Cell Systems 17 (6): 101574.

Goswami, Garv, David Xu, and Hwi Joo Park. 2026. “Contrastive Alignment of Expression and Copy Number Highlights Dosage-Insensitive Genes in Cancer.” In bioRxiv. BioRxiv, March 3. 10.64898/2026.03.01.708901.

Gyorgypal, Aron, Antash Chaturvedi, Viki Chopda, Haoran Zhang, and Shishir P. S. Chundawat. 2025. “Evaluating the Impact of Media and Feed Combinations on CHO Cell Culture Performance and Monoclonal Antibody (trastuzumab) Production.” Cytotechnology 77 (1): 40.

Hamaya, Yamato, Ayako Suzuki, Yutaka Suzuki, Katsuya Tsuchihara, and Riu Yamashita. 2023. “Classification and Characterization of Alternative Promoters in 26 Lung Adenocarcinoma Cell Lines.” Japanese Journal of Clinical Oncology 53 (2): 97–104.

Heffner, Kelley, Deniz Baycin Hizal, Natalia I. Majewska, et al. 2020. “Expanded Chinese Hamster Organ and Cell Line Proteomics Profiling Reveals Tissue-Specific Functionalities.” Scientific Reports 10 (1): 15841.

Hernandez Gifford, J. A. 2015. “The Role of WNT Signaling in Adult Ovarian Folliculogenesis.” Reproduction 150 (4): R137–48.

Hernandez, Inmaculada, Heena Dhiman, Gerald Klanert, et al. 2019. “Epigenetic Regulation of Gene Expression in Chinese Hamster Ovary Cells in Response to the Changing Environment of a Batch Culture.” Biotechnology and Bioengineering 116 (3): 677–692.

Hoffman, Gabriel E., and Eric E. Schadt. 2016. “variancePartition: Interpreting Drivers of Variation in Complex Gene Expression Studies.” BMC Bioinformatics 17 (1): 483.

Holgersson, Jan, and Jonas Löfling. 2006. “Glycosyltransferases Involved in Type 1 Chain and Lewis Antigen Biosynthesis Exhibit Glycan and Core Chain Specificity.” Glycobiology 16 (7): 584–593.

Ho, Wan-Ling, Wen-Ming Hsu, Min-Chuan Huang, Kenji Kadomatsu, and Akira Nakagawara. 2016. “Protein Glycosylation in Cancers and Its Potential Therapeutic Applications in Neuroblastoma.” Journal of Hematology & Oncology 9 (1): 100.

Huhn, Steve, Meiping Chang, Amit Kumar, et al. 2022. “Chromosomal Instability Drives Convergent and Divergent Evolution toward Advantageous Inherited Traits in Mammalian CHO Bioproduction Lineages.” iScience 25 (4): 104074.

Hu, Lixia, Shanshan Liu, Lisha Xia, et al. 2022. “CHO Cathepsin B Identified as the Protease Responsible for a Target Bispecific Antibody Fragmentation.” Protein Expression and Purification 199 (106144): 106144.

Hu, Zhilan, Donglin Guo, Shirley S. M. Yip, et al. 2013. “Chinese Hamster Ovary K1 Host Cell Enables Stable Cell Line Development for Antibody Molecules Which Are Difficult to Express in DUXB11-Derived Dihydrofolate Reductase Deficient Host Cell.” Biotechnology Progress (111 RIVER ST, HOBOKEN 07030-5774, NJ) 29 (4): 980–985.

Kaas, Christian Schrøder, Claus Kristensen, Michael J. Betenbaugh, and Mikael Rørdam Andersen. 2015. “Sequencing the CHO DXB11 Genome Reveals Regional Variations in Genomic Stability and Haploidy.” BMC Genomics 16 (1): 160.

Kim, Daehwan, Joseph M. Paggi, Chanhee Park, Christopher Bennett, and Steven L. Salzberg. 2019. “Graph-Based Genome Alignment and Genotyping with HISAT2 and HISAT-Genotype.” Nature Biotechnology 37 (8): 907–915.

Lakshmanan, Meiyappan, Yee Jiun Kok, Alison P. Lee, et al. 2019. “Multi-Omics Profiling of CHO Parental Hosts Reveals Cell Line-Specific Variations in Bioprocessing Traits.” Biotechnology and Bioengineering 116 (9): 2117–2129.

Lewis, Nathan E., Xin Liu, Yuxiang Li, et al. 2013. “Genomic Landscapes of Chinese Hamster Ovary Cell Lines as Revealed by the Cricetulus Griseus Draft Genome.” Nature Biotechnology 31 (8): 759–765.

Liao, Yang, Gordon K. Smyth, and Wei Shi. 2014. “featureCounts: An Efficient General Purpose Program for Assigning Sequence Reads to Genomic Features.” Bioinformatics (Oxford, England) 30 (7): 923–930.

Louie, Salina, Amy Heidersbach, Noelia Blanco, et al. 2020. “Endothelial Intercellular Cell Adhesion Molecule 1 Contributes to Cell Aggregate Formation in CHO Cells Cultured in Serum-Free Media.” Biotechnology Progress 36 (3): e2951.

Love, Michael I., Wolfgang Huber, and Simon Anders. 2014. “Moderated Estimation of Fold Change and Dispersion for RNA-Seq Data with DESeq2.” Genome Biology 15 (12): 550.

Malm, Magdalena, Rasool Saghaleyni, Magnus Lundqvist, et al. 2020. “Evolution from Adherent to Suspension: Systems Biology of HEK293 Cell Line Development.” Scientific Reports 10 (1): 18996.

Martin, Marcel. 2011. “Cutadapt Removes Adapter Sequences from High-Throughput Sequencing Reads.” EMBnet.journal 17 (1): 10.

Masson, Helen, Jasmine Tat, Pablo Di Giusto, et al. 2025. “A Reconstruction of the Mammalian Secretory Pathway Identifies Mechanisms Regulating Antibody Production.” Cell Systems 16 (12): 101453.

Ma, Xiaonan, Ling Zhang, Luming Zhang, et al. 2020. “Validation and Identification of Reference Genes in Chinese Hamster Ovary Cells for Fc-Fusion Protein Production.” Experimental Biology and Medicine (Maywood, N.J.) 245 (8): 690–702.

McInnes, Leland, John Healy, Nathaniel Saul, and Lukas Großberger. 2018. “UMAP: Uniform Manifold Approximation and Projection.” Journal of Open Source Software 3 (29): 861.

Moosmann, P., O. Georgiev, H. J. Thiesen, M. Hagmann, and W. Schaffner. 1997. “Silencing of RNA Polymerases II and III-Dependent Transcription by the KRAB Protein Domain of KOX1, a Krüppel-Type Zinc Finger Factor.” Biological Chemistry 378 (7): 669–677.

Rask, K., A. Nilsson, M. Brännström, et al. 2003. “Wnt-Signalling Pathway in Ovarian Epithelial Tumours: Increased Expression of Beta-Catenin and GSK3beta.” British Journal of Cancer 89 (7): 1298–1304.

Reinhart, David, Lukas Damjanovic, Christian Kaisermayer, et al. 2019. “Bioprocessing of Recombinant CHO-K1, CHO-DG44, and CHO-S: CHO Expression Hosts Favor Either mAb Production or Biomass Synthesis.” Biotechnology Journal 14 (3): e1700686.

Reinhart, David, Lukas Damjanovic, Christian Kaisermayer, and Renate Kunert. 2015. “Benchmarking of Commercially Available CHO Cell Culture Media for Antibody Production.” Applied Microbiology and Biotechnology 99 (11): 4645–4657.

Riedl, Markus, Nicolas Marx, and Nicole Borth. 2026. “Global Structure and Determinants of Transcriptomic Adaptation in CHO Cells.” New Biotechnology 96 (September): 32–43.

Riedl, Markus, Caterina Ruggeri, Nicolas Marx, and Nicole Borth. 2025. “Fantastic Genes and Where to Find Them Expressed in CHO.” Computational and Structural Biotechnology Journal 27 (April): 1407–1415.

Ritchie, Matthew E., Belinda Phipson, Di Wu, et al. 2015. “Limma Powers Differential Expression Analyses for RNA-Sequencing and Microarray Studies.” Nucleic Acids Research 43 (7): e47.

Robinson, Mark D., Davis J. McCarthy, and Gordon K. Smyth. 2010. “edgeR: A Bioconductor Package for Differential Expression Analysis of Digital Gene Expression Data.” Bioinformatics (Oxford, England) 26 (1): 139–140.

Robinson, Mark D., and Alicia Oshlack. 2010. “A Scaling Normalization Method for Differential Expression Analysis of RNA-Seq Data.” Genome Biology 11 (3): R25.

Rodrigues, Maria Elisa, Ana Rita Costa, Mariana Henriques, Joana Azeredo, and Rosário Oliveira. 2012. “Comparison of Commercial Serum-Free Media for CHO-K1 Cell Growth and Monoclonal Antibody Production.” International Journal of Pharmaceutics 437 (1-2): 303–305.

Ronemus, Michael, Daniel Bradford, Zachary Laster, and Siran Li. 2025. “Exploring Genome-Transcriptome Correlations in Cancer.” Biochemical Society Transactions 53 (1): BST20240108.

Rupp, Oliver, Jennifer Becker, Karina Brinkrolf, et al. 2014. “Construction of a Public CHO Cell Line Transcript Database Using Versatile Bioinformatics Analysis Pipelines.” PloS One 9 (1): e85568.

Rupp, Oliver, Madolyn L. MacDonald, Shangzhong Li, et al. 2018. “A Reference Genome of the Chinese Hamster Based on a Hybrid Assembly Strategy.” Biotechnology and Bioengineering 115 (8): 2087–2100.

Sedighikamal, Hossein, Reza Karimi Mostofi, Alireza Sattarzadeh, Mansour Shahbazi, and Hossein Aghazadeh. 2022. “Comparative Study of Commercial Media to Improve GMP Manufacturing of Recombinant Human Interferon β-1a by CHO Cells in Perfusion Bioreactor.” Cytotechnology 74 (6): 669–680.

Shao, Xin, Ning Lv, Jie Liao, et al. 2019. “Copy Number Variation Is Highly Correlated with Differential Gene Expression: A Pan-Cancer Study.” BMC Medical Genetics 20 (1): 175.

Shepherd, L, Morgan, M. 2025. AnnotationHub. Bioconductor. 10.18129/B9.BIOC.ANNOTATIONHUB.

Singh, Ankita, Helene F. Kildegaard, and Mikael R. Andersen. 2018. “An Online Compendium of CHO RNA-Seq Data Allows Identification of CHO Cell Line-Specific Transcriptomic Signatures.” Biotechnology Journal 13 (10): e1800070.

Széliová, Diana, David E. Ruckerbauer, Sarah N. Galleguillos, et al. 2020. “What CHO Is Made of: Variations in the Biomass Composition of Chinese Hamster Ovary Cell Lines.” Metabolic Engineering 61 (September): 288–300.

Takahara, Terunao, Yuna Amemiya, Risa Sugiyama, Masatoshi Maki, and Hideki Shibata. 2020. “Amino Acid-Dependent Control of mTORC1 Signaling: A Variety of Regulatory Modes.” Journal of Biomedical Science 27 (1): 87.

Tejeda-Muñoz, Nydia, and Kuo-Ching Mei. 2024. “Wnt Signaling in Cell Adhesion, Development, and Colon Cancer.” IUBMB Life 76 (7): 383–396.

Uhlén, Mathias, Linn Fagerberg, Björn M. Hallström, et al. 2015. “Proteomics. Tissue-Based Map of the Human Proteome.” Science (New York, N.Y.) 347 (6220): 1260419.

Vishwanathan, Nandita, Arpan Bandyopadhyay, Hsu-Yuan Fu, Kathryn C. Johnson, Nathan M. Springer, and Wei-Shou Hu. 2017. “A Comparative Genomic Hybridization Approach to Study Gene Copy Number Variations among Chinese Hamster Cell Lines: Gene Copy Number Variation in Cho Cell Lines.” Biotechnology and Bioengineering 114 (8): 1903–1908.

Vishwanathan, Nandita, Andrew Yongky, Kathryn C. Johnson, et al. 2015. “Global Insights into the Chinese Hamster and CHO Cell Transcriptomes: Chinese Hamster and CHO Cell Transcriptomes.” Biotechnology and Bioengineering 112 (5): 965–976.

Wang, Shiyan, Chi Chun Wong, Yanquan Zhang, et al. 2021. “ZNF545 Loss Promotes Ribosome Biogenesis and Protein Translation to Initiate Colorectal Tumorigenesis in Mice.” Oncogene 40 (48): 6590–6600.

Wurm, Florian. 2013. “CHO Quasispecies—implications for Manufacturing Processes.” Processes (Basel, Switzerland) 1 (3): 296–311.

Wu, Tianzhi, Erqiang Hu, Shuangbin Xu, et al. 2021. “clusterProfiler 4.0: A Universal Enrichment Tool for Interpreting Omics Data.” Innovation (Cambridge (Mass.)) 2 (3): 100141.

Xu, Xun, Harish Nagarajan, Nathan E. Lewis, et al. 2011. “The Genomic Sequence of the Chinese Hamster Ovary (CHO)-K1 Cell Line.” Nature Biotechnology 29 (8): 735–741.

Yamano-Adachi, Noriko, Yuto Nakanishi, Wataru Tanaka, et al. 2023. “Artificial Induction of Chromosome Aneuploidy in CHO Cells Alters Their Function as Host Cells.” Biotechnology and Bioengineering 120 (3): 659–673.

Yang, Zhang, Adnan Halim, Yoshiki Narimatsu, et al. 2014. “The GalNAc-Type O-Glycoproteome of CHO Cells Characterized by the SimpleCell Strategy.” Molecular & Cellular Proteomics 13 (12): 3224–3235.

Yatipanthalawa, Bhagya S., Yih Yean Lee, Sally L. Gras, and Gregory J. O. Martin. 2026. “Amino Acid Metabolism, Demand and Supply in Chinese Hamster Ovary Cell Culture - A Comprehensive Literature Review.” Biotechnology Advances 89 (108866): 108866.

Yu, Michael, Kevin Qin, Jiaming Fan, et al. 2024. “The Evolving Roles of Wnt Signaling in Stem Cell Proliferation and Differentiation, the Development of Human Diseases, and Therapeutic Opportunities.” Genes & Diseases 11 (3): 101026.

